# Regulation of lipid saturation without sensing membrane fluidity

**DOI:** 10.1101/706556

**Authors:** Stephanie Ballweg, Erdinc Sezgin, Dorith Wunnicke, Inga Hänelt, Robert Ernst

## Abstract

Cells maintain membrane fluidity by regulating lipid saturation, but the molecular mechanisms of this homeoviscous adaptation remain poorly understood. Here, we have reconstituted the core machinery for sensing and regulating lipid saturation in baker’s yeast to directly characterize its response to defined membrane environments. Using spectroscopic techniques and *in vitro* ubiquitylation, we uncover a unique sensitivity of the transcriptional regulator Mga2 to the abundance, position, and configuration of double bonds in lipid acyl chains and provide unprecedented insight into the molecular rules of membrane adaptivity. Our data challenge the prevailing hypothesis that membrane viscosity serves as the measured variable for regulating lipid saturation. Rather, we show that the signaling output of Mga2 correlates with the size of a single sensor residue in the transmembrane helix, which senses the lateral pressure and/or compressibility profile in a defined region of the membrane. Our findings suggest that membrane property sensors have evolved remarkable sensitivities to highly specific aspects of membrane structure and dynamics, thus paving the way toward the development of genetically encoded reporters for such membrane properties in the future.

## Introduction

Cellular membranes are complex assemblies of proteins and lipids, which collectively determine physical bilayer properties such as membrane viscosity, permeability, and the lateral pressure profile^1–4^. The acyl chain composition of membrane lipids is an important determinant of membrane viscosity and tightly controlled in bacteria^5–7^, fungi^8,9^, worms^10,11^, flies^12^, and vertebrates^13,14^. Saturated lipid acyl chains tend to form non-fluid, tightly packed gel phases, while unsaturated lipid acyl chains fluidize the bilayer. Poikilothermic organisms that cannot control their body temperature must adjust their lipid composition during cold stress to maintain membrane functions– a phenomenon referred to as the homeoviscous adaptation^15–17^. Despite recent advances in identifying candidate sensory, it remains largely unknown how these sensors work on the molecular scale and how they are coordinated for maintaining a physicochemical membrane homeostasis^20,21^. The fact that most, if not all, membrane properties are interdependent is a key challenge for this emerging field. How do cells, for example, balance the need for maintaining membrane viscosity with the need to maintain organelle-specific lateral pressure profiles^22^? In fact, perturbation of membrane viscosity by genetically targeting fatty acid metabolism leads to complex changes throughout the entire lipidome impacting on other bilayer properties and causing endoplasmic reticulum (ER) stress and disruption of its normal architecture^19,23,24^. Clearly, we are lacking a unifying theory that could accurately predict the properties of a membrane when the composition is known: Each component of a complex biological membrane contributes to the collective, physicochemical properties in a non-ideal, non-additive, and non-linear fashion^3,25^. As important step towards a unifying membrane theory, we need to identify a set of membrane properties, which are minimally correlated and sufficient to uniquely describe the state of a bilayer. Characterizing naturally occurring membrane property sensors, which may exhibit highly specialized sensitivities to specific membrane properties, holds promise to better understand how cells prioritize the maintenance of such orthogonal membrane properties^21^.

Eukaryotic cells use sensor proteins possessing refined mechanisms to monitor physicochemical properties of their organellar membranes and to adjust lipid metabolism during stress, metabolic adaptation, and development^10,24,26–31^. These sensor proteins can be categorized into three classes, based on topological considerations^20,21^: Class I sensors interrogate surface properties of cellular membranes, such as the surface charge and molecular packing density as reported for amphipathic lipid packing sensor (ALPS) motif containing proteins and other amphipathic helix containing proteins^32^. Class II sensors perturb and interrogate the hydrophobic core of the bilayer and have been implicated in the regulation of lipid saturation. Class III sensors are transmembrane proteins acting across the bilayer by locally squeezing, stretching, and/or bending the membrane to challenge selective properties such as thickness or bending rigidity^20,21^.

The prototypical class II sensor Mga2 is crucial for the regulation of membrane viscosity in the baker’s yeast^9,26^ (Figure 1A). Its single transmembrane helix (TMH) senses a physicochemical signal in the ER membrane to control a homeostatic response that adjusts membrane lipid saturation via the essential fatty acid *cis*-Δ*9*-desaturase Ole1^33–35^. Increased lipid saturation triggers the ubiquitylation of three lysine residues in the cytosolic, juxtamembrane region of Mga2 by the E3 ubiquitin ligase Rsp5^36^. This ubiquitylation serves as a signal for the proteasome-dependent processing of the membrane-bound Mga2 precursor (P120) and the release of a transcriptionally active P90 fragment, which upregulates *OLE1* expression ^37^ (Figure 1A). This regulated, ubiquitin/proteasome dependent processing resembles the pathway of ER-associated degradation^38^ and was first described for Spt23, a close structural and functional homologue of Mga2^39^. Because Ole1 is the only source for the *de novo* biosynthesis of unsaturated fatty acids, its tight regulation via Mga2 is essential for maintaining membrane fluidity in this poikilotherm^9,35^.

**Figure 1:**
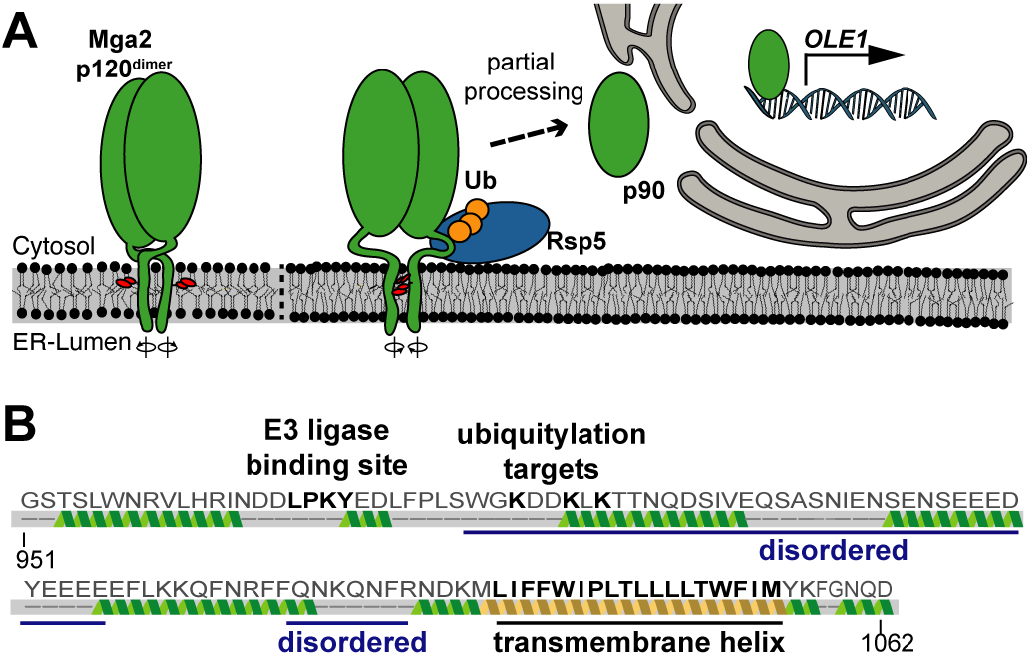
The activation of Mga2 is controlled by the ER membrane composition. **(A)** Model of the OLE pathway: the transcription factor Mga2 forms inactive dimers in the ER membrane (Mga2 p120^dimer^) with highly dynamic TMHs exploring alternative rotational orientations. Loose lipid packing (left) caused by unsaturated lipids stabilizes conformations with two sensory tryptophan residues (W1042; red) pointing away from the dimer interface toward the lipid environment. Tight lipid packing (right) stabilizes alternative rotational conformations with the sensory tryptophans facing each other in the dimer interface (right). The E3 ubiquitin ligase Rsp5 is required to ubiquitylate (Ub) Mga2, thereby facilitating the proteolytic processing by the proteasome and the release of transcriptionally active Mga2 (p90). **(B)** Secondary structure prediction of the juxtamembrane and transmembrane region (residue 951-1062) of Mga2 using Phyre2^61^.

Molecular dynamics (MD) simulations have revealed a remarkable conformational flexibility of the Mga2 transmembrane region^26^. The TMHs of Mga2 dimerize and rotate against each other, thus forming an ensemble of dimerization interfaces. Importantly, the population of these alternative configurations is affected by the membrane lipid environment: Higher proportions of saturated lipid acyl chains stabilize a configuration, in which two tryptophan residues (W1042) point towards the dimer interface, whereas higher proportions of unsaturated lipid acyl chains favor a conformation where these residues point away from another and toward the lipid environment^9,26^. Based on the remarkable correspondence with genetic and biophysical data, we proposed that this membrane-dependent, structural dynamics of the TMHs are coupled to the ubiquitylation and activation of Mga2^26^. However, it remained uncertain if the reported, relatively subtle changes in the population of short-lived, rotational conformations are sufficient to control a robust cellular response. How can the processing of Mga2 be blocked by an increased proportion of unsaturated lipids in the membrane, if the sensory TMHs still explore their entire conformational space? How is the ‘noisy’ signal from the TMH propagated via disordered regions to the site of ubiquitylation in the juxtamembrane region (Figure 1B)?

As an important step toward answering these questions, we have designed and isolated a second-generation, minimal sensor construct based on Mga2. It senses the membrane environment and acquires, depending on the membrane lipid composition, a poly-ubiquitylation label as a signal for its activation via proteasomal processing. After reconstituting this sense-and-response construct in liposomes with defined lipid compositions, we demonstrate a remarkable sensitivity of Mga2 to specific changes in the bilayer composition. We provide compelling evidence for functional coupling between the TMH and the site of ubiquitylation using electron paramagnetic resonance (EPR) and Förster-resonance energy transfer (FRET). Strikingly, our data rule out the hypothesis that Mga2 acts as a sensor for membrane viscosity/fluidity. Instead, we propose based on our findings that Mga2 senses a small portion of the lateral pressure and/or lateral compressibility profile via the sensory tryptophan (W1042)^26^ within the hydrophobic core of the membrane. Thus, our mechanistic analysis of the membrane lipid saturation sensor Mga2 challenges the common view of membrane viscosity as the critical measured variable in membrane biology.

## Results

### A minimal sense-and-response construct reports on membrane lipid saturation

We proposed that Mga2 uses a rotation-based mechanism to sense membrane lipid saturation^26^ (Figure 1A). However, the sensory TMHs of Mga2 are separated from the site of ubiquitylation by a predicted disordered loop and ~50 amino acids (Figure 1B), thereby posing a question of their functional coupling. How can the conformational dynamics of the TMHs control the ubiquitylation of Mga2 in the juxtamembrane region? In order to study the coupling of sensing and ubiquitylation *in vitro*, we have generated a minimal sense-and-response construct (^ZIP-MBP^Mga2^950-1062^) comprising an N-terminal leucine-zipper (ZIP) derived from the transcription factor GCN4, the maltose binding protein (MBP), the juxtamembrane region (950-1036) and the TMH (1037-1058) of Mga2 (950-1062) (Figure 2A). The N-terminal zipper mimics the IPT (Ig-like, plexin, transcription factor)-domain of full-length Mga2 and stabilizes a homo-dimeric state^40^, while the MBP was used as a purification and solubility tag ^26^. The juxtamembrane domain of Mga2 comprises the LPKY motif (Mga2^958-961^) for recruiting the E3 ubiquitin ligase Rsp5^41^, three lysine residues K^980^, K^983^ and K^985^ ubiquitylated *in vivo*^36^, and the disordered region linking these motifs to the TMH (Figure 2A). The construct was recombinantly produced and isolated in the presence of Octyl-β-D-glucopyranoside (OG) using an amylose-coupled affinity matrix and size-exclusion chromatography (SEC) (Figure 2B, S1A). Expectedly, the N-terminal zipper stabilizes a dimeric form of the sense-and-response construct and supports, at increased concentrations, the formation of higher oligomeric forms as suggested by SEC experiments that also included a zipper-less variant (^MBP^Mga2^950-1062^) as a control (Figure 2C, S1B, C). We reconstituted the sense-and-response construct in liposomes at molar protein-to-lipid ratios between 1:5,000 – 1:15,000 and detected no sign of protein aggregation in our preparations using sucrose-density gradient centrifugations (Figure S1D).

**Figure 2:**
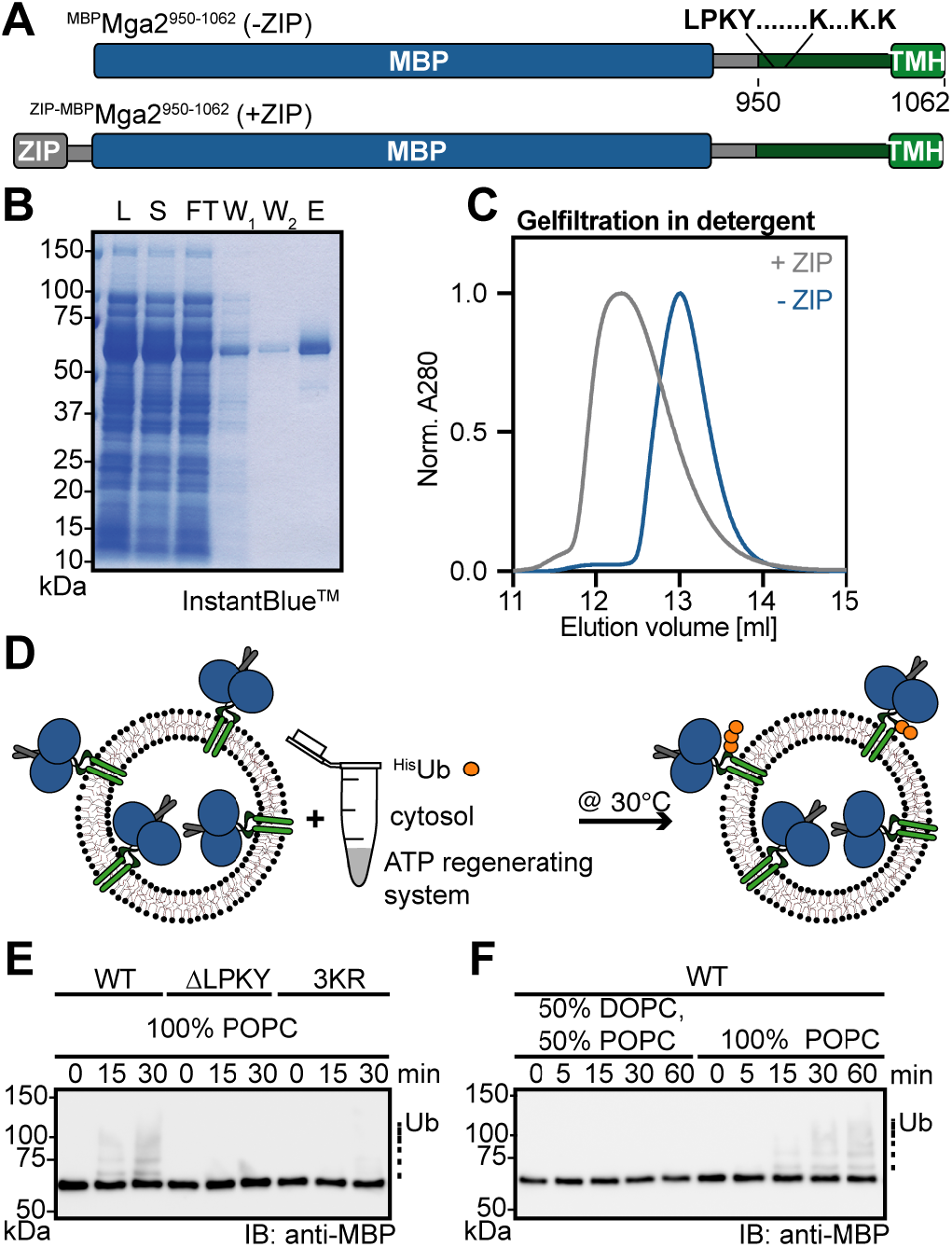
An *in vitro* sense-and-response system for membrane lipid saturation. (**A**) Schematic representation of the sense-and-response constructs. The fusion proteins are composed of the maltose-binding protein (MBP, blue) and Mga2^950-1062^, encompassing the Rsp5 binding site (LPKY), three lysine residues targeted for ubiquitylation (K^980^, K^983^ and K^985^), a predicted disordered juxta-membrane region, and the C-terminal TMH (green). An optional, N-terminal leucine zipper derived from Gcn4 (grey, residues 249-281) was used to support dimerization. **(B)** Isolation of the zipped sense-and-response construct by affinity purification. 0.1 OD units of the lysate (L), soluble (S), flow-through (FT), and two wash fractions (W_1,2_), as well as 1 *μ*g of the eluate were subjected to SDS-PAGE followed by InstantBlue^TM^ staining. The protein was further purified by preparative SEC (FigS1A). (**C**) 100 *μ*g in 100 *μ*l of the purified sense-and-response constructs with (+ZIP) and without zipper (−ZIP) were loaded on a Superdex 200 10/300 Increase column (void volume 8.8 ml). (**D**) Schematic representation of the *in vitro* ubiquitylation assay. Proteoliposomes containing ^ZIP-MBP^Mga2^950-1062^ were mixed with ^8xHis^Ubiquitin (^His^Ub), an ATP-regenerating system, and cytosol prepared from wildtype yeasts to facilitate Mga2 ubiquitylation at 30°C. (**E**) The reaction was performed with the ^ZIP-MBP^Mga2^950-1062^ wildtype (WT) sense-and-response construct, a variant lacking the Rsp5 binding site (ΔLPKY), and a variant with three substitutions of the lysine residues K^980^, K^983^, and K^985^ to arginine (3KR), thereby removing the targets of Rsp5-dependent *in vivo* ubiquitylation. The constructs were reconstituted in liposomes composed of 100 mol% POPC at a protein-to-lipid ratio of 1:5,000. After indicated times, the reactions were stopped using sample buffer and subjected to SDS-PAGE. For analysis, an immunoblot using anti-MBP antibodies was performed. (**F**) Ubiquitylation reactions were performed as in (E) with the WT sense-and-response construct reconstituted in the indicated lipid environments at a molar protein-to-lipid ratio of 1:5,000.

We then tested if the sense-and-response construct could be ubiquitylated *in vitro* and adapted a strategy established for the ubiquitylation of substrates of the ER-associated degradation (ERAD) machinery^42^. We incubated the proteoliposomes with an ATP-regenerating system, purified ^8xHis^ubiquitin, and yeast cytosol containing enzymes required to mediate the ubiquitylation reaction (Figure 2D). Subsequent immunoblot analyses revealed a time-dependent ubiquitylation of the sense-and-response construct, which became apparent as a ladder of MBP-positive signals (Figure 2E). Control experiments validated the specificity of the ubiquitylation reaction: No ubiquitylation was observed, when the Rsp5-binding site (ΔLPKY) was deleted from the sense-and-response construct (Figure 2E). Furthermore, despite the presence of 50 lysine residues in the entire construct, the substitution of the three lysine residues (3KR) targeted by Rsp5 *in vivo* ^36^ was sufficient to prevent the ubiquitylation (Figure 2E). We conclude that the *in vitro* ubiquitylation assay is specific and that the conformational dynamics in the juxtamembrane region is likely to reflect the structural dynamics found in full-length Mga2 protein. Most importantly, this newly established *in vitro* system also allowed us to test the hypothesis of functional coupling between the sensory TMHs and protein ubiquitylation.

We reconstituted the sense-and-response construct in two distinct membrane environments based on a phosphatidylcholine (PC) matrix but differing in their lipid acyl chain composition. One membrane environment contained 50% unsaturated 18:1 and 50% saturated 16:0 acyl chains (100 mol% POPC(16:0/18:1)), while the other was less saturated and contained 75% unsaturated 18:1 and 25 saturated 16:0 acyl chains (50 mol% DOPC(18:1/18:1), 50 mol% POPC) (Figure 2F). Notably, this degree of lipid saturation is in the range of the naturally occurring acyl chain compositions reported for baker’s yeast cultivated in different conditions^8,23,43,44^. The sense-and-response construct was efficiently ubiquitylated in the more saturated membrane environment (evidenced by the appearance of bands with decreased electrophoretic mobility), but not in the unsaturated one (Figure 2F, S1E). This observation highlights the remarkable sensitivity of class II membrane property sensor and provides strong evidence for a functional coupling between the TMHs and the site of ubiquitylation.

### An *in vitro* strategy to reconstitute membrane lipid sensing

In order to detect changes of the conformational dynamics in the juxtamembrane region, we established an *in vitro* FRET assay. We hypothesized that the average distance between the binding site of the E3 ligase Rsp5 (LPKY) and a lysine residue targeted by Rsp5 may be affected by changes in the membrane lipid environment. We thus generated a donor construct labeled with Atto488 at the position of a target-lysine (K983^D^) and an acceptor construct labeled with Atto590 within the Rsp5 recognition site (K969^A^) (Förster radius of 59 Å) (Figure 3A). Notably, the required amino acid substitutions to cysteine at the positions of labeling did not interfere with the activation of full-length Mga2 *in vivo* (Figure S2A). The individually isolated donor (K983^D^) and acceptor (K969^A^) constructs exhibited only negligible fluorescence emission at 614 nm in detergent solution upon donor excitation at 488 nm (Figure 3B). However, a significant emission at 614 nm (from here on referred to as FRET signal) was detectable upon mixing the donor and acceptor constructs (K983^D^+K969^A^) (Figure 3B). Notably, a direct excitation of the acceptor at 590 nm (Figure S2B) resulted in equal fluorescence intensities at 614 nm for both K983^D^+K969^A^ and K969^A,only^ samples, but no emission for the K983^D,only^ sample. The normalized FRET signal of the K983^D^+K969^A^ reporter was concentration-dependent in detergent solution (Figure 3C), thereby suggesting a dynamic equilibrium between monomeric and oligomeric species (presumably dimers) of the labeled sense-and-response construct. To validate this interpretation and to rule out the possibility that the FRET signal was predominantly caused by FRET between stable K983^D^-K983^D^ and K969^A^-K969^A^ dimers bumping into each other, we performed competition experiments. We found that the ratiometric FRET efficiency of the K983^D^+K969^A^ reporter was substantially reduced upon titrating it with an unlabeled sense-and-response construct containing an N-terminal leucine-zipper (Figure 3D). However, it remained unaffected upon titration with an unlabeled construct lacking a zipper (Figure 3D). This indicates (i) that the zipper centrally contributes to the stability of the dimer, (ii) that individual protomers readily exchange in detergent solution, and (iii) that the FRET signal is mainly due to K983^D^-K969^A^ heterooligomers. In fact, additional titration experiments with the K969^A^ acceptor revealed that the observed FRET efficiency is a linear function of the molar fraction of the acceptor (Figure S2C,D), thereby indicating that the FRET signal is indeed caused by dimers^45^.

**Figure 3:**
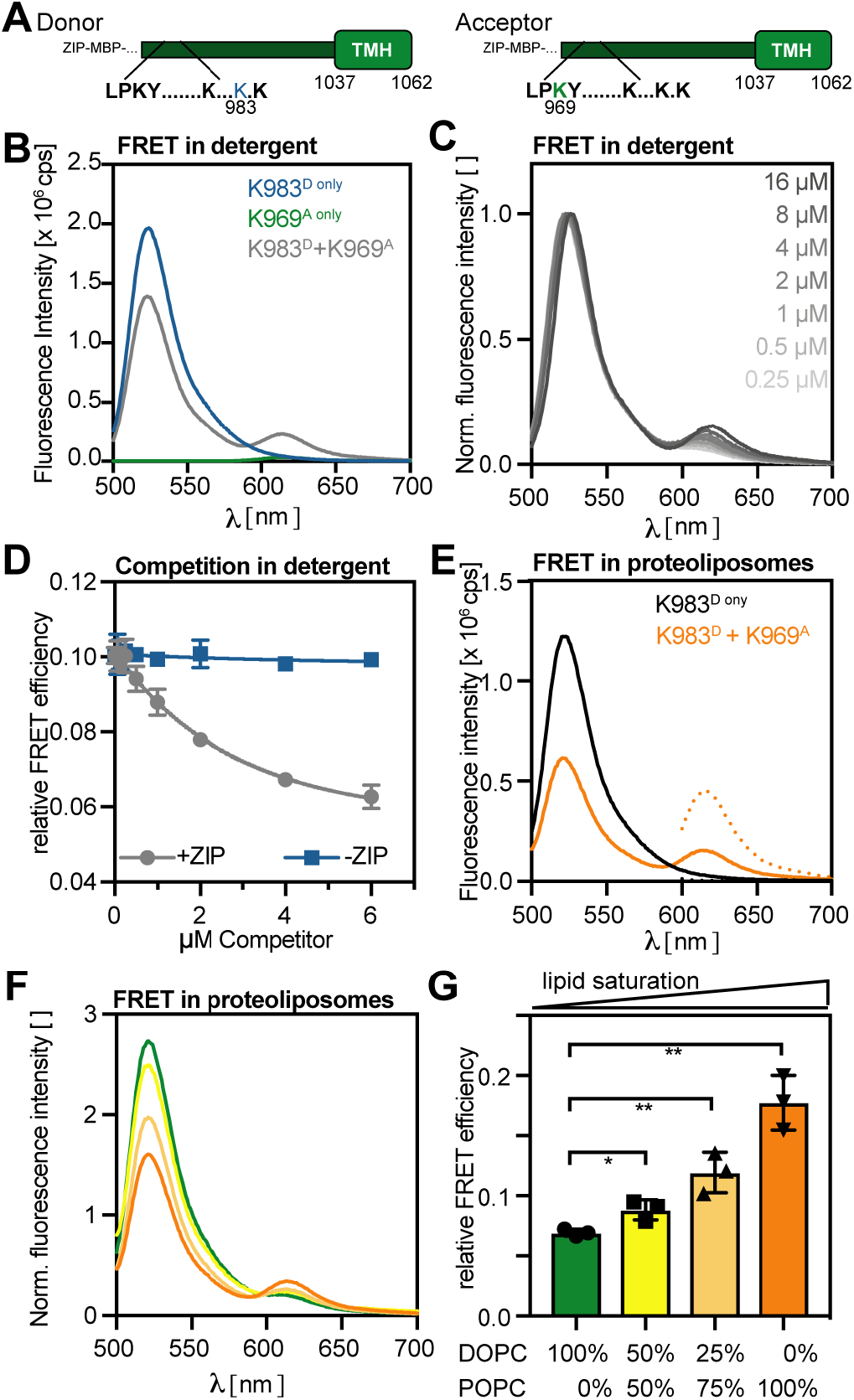
FRET reveals membrane-controlled, conformational changes in the sense- and-response construct. **(A)** Schematic representation of the donor and acceptor construct. The Atto488 donor was installed at the position K^983^ via a cysteine mutant, thereby substituting a residue that is ubiquitylated by Rsp5 *in vivo*. The Atto590 acceptor was installed via a K969C mutation in the Rsp5 binding site. **(B)** Fluorescence emission spectra reveal energy transfer between donor and acceptor in detergent solution. 2 *μ*M of each construct was used to record fluorescence emission spectra (ex: 488 nm, em: 500-700 nm) of the donor (K983^D only^), acceptor (K969^A only^), and the combined (K983^D^+K969^A^) FRET pair. **(C)** Fluorescence emission spectra were recorded for serial dilutions of the donor/acceptor pair in detergent as in (B). The spectra were normalized to maximal fluorescence intensity at the donor emission. **(D)** Zipped donor (2 *μ*M) and acceptor (2 *μ*M) pairs were premixed in detergent solution for 10 min to allow for protomer exchange and full equilibration before unlabeled competitor constructs with (+ZIP) and without zipper (-ZIP) were added to the indicated concentration. Fluorescence spectra were recorded as in (B). The relative FRET efficiency was determined from the ratio of the donor/acceptor intensities and plotted as the mean ± SD from two independent experiments. (E) Fluorescence emission spectra indicate energy transfer within the membrane-reconstituted sense-and-response construct. The donor construct was premixed either with unlabeled (K983^D only^) or labeled acceptor construct (K983^D^+K969^A^) prior to a reconstitution in POPC liposomes at a protein-to-lipid ratio of 1:8,000. Fluorescence emission spectra (em500-700 nm) from donor excited (ex: 488 nm; solid line) and acceptor excited (ex: 590 nm; dotted line) samples. **(F)** Donor (K983^D^) and acceptor (K969^A^) were premixed and incubated in detergent solution at a molar ratio of 1:1 and used for reconstitution in liposomes with indicated lipid compositions. Fluorescence emission spectra were recorded as in (E) and normalized to the maximal acceptor emission after direct acceptor excitation (ex: 590 nm). **(G)** The relative FRET efficiency was as in (F) and plotted as the mean ± SD of three independent measurements. A two-tailed, unpaired t-test was performed to test for statistical significance (*p<0.01, **p<0.001).

Next, we studied the structural dynamics of the sense-and-response construct in liposomes using the FRET reporter. To this end, we reconstituted K983^D,only^ and the pre-mixed K983^D^+K969^A^ pair in liposomes of defined lipid compositions and recorded fluorescence spectra (Figure 3E-G). We used a low protein-to-lipid ratio of 1:8,000 in these experiments to minimize the contribution of unspecific proximity FRET to the overall signal^45^. We observed a significant FRET signal for the K983^D^-K969^A^ reporter reconstituted in a POPC bilayer (Figure 3E) evidenced by a decreased donor fluorescence and an increased acceptor emission at 614 nm compared to the K983^D,only^ sample (Figure 3E). Using this FRET assay, we then studied the impact of the lipid acyl chain composition on the structural dynamics of the juxtamembrane region. The lowest FRET efficiency was observed in a DOPC bilayer containing 100% unsaturated acyl chains (Figure 3F,G). At higher proportions of saturated lipid acyl chains in the bilayer the FRET efficiency increased. These data demonstrate that the acyl chain composition in the hydrophobic core of the membrane imposes structural changes to regions outside the membrane, which have been implicated in signal propagation^36,41^. Our data establish an intricate functional and structural coupling between the TMH regions and the sites of ubiquitylation.

### The Mga2-based sense-and-response construct does not report on membrane viscosity

The remarkable sensitivity of Mga2 to lipid saturation raises the question if this is based on sensing membrane viscosity. In order to test this hypothesis, we first measured the diffusion coefficients of fluorescent lipid analogues (0.01 mol% Atto488-DPPE (Figure 4A) and 0.01 mol% Abberior Star Red-PEG Cholesterol) (Figure S3A) in giant unilaminar vesicles with different lipid compositions via confocal point fluorescence correlation spectroscopy (FCS). Expectedly, the membrane viscosity increases slightly with the proportion of saturated lipid acyl chains (from 0% saturated acyl chains for DOPC to 50% for POPC) as evidenced by decreasing diffusion coefficients of the labeled lipids (Figure 4A, S3A). Previous reports have identified a central contribution of phosphatidylethanolamine (PE) to membrane viscosity in cells^46^. Consistently, PE increases the membrane viscosity in model membranes: A lipid bilayer composed of 60 mol% PC and 40 mol% PE with 25% saturated lipid acyl chains is significantly more viscous than other bilayers composed of only PC with 0%, 25%, or even 50% saturated acyl chains (Figure 4A). We also studied these lipid compositions by C-laurdan spectroscopy, which reports on water penetration into the lipid bilayer^47^. A low degree of water penetration increases the generalized polarization (GP) of C-laurdan and indicates tighter lipid packing. For the investigated set of lipids, the membrane viscosity correlated with the respective degree of lipid packing (Figure 4 A, B).

**Figure 4:**
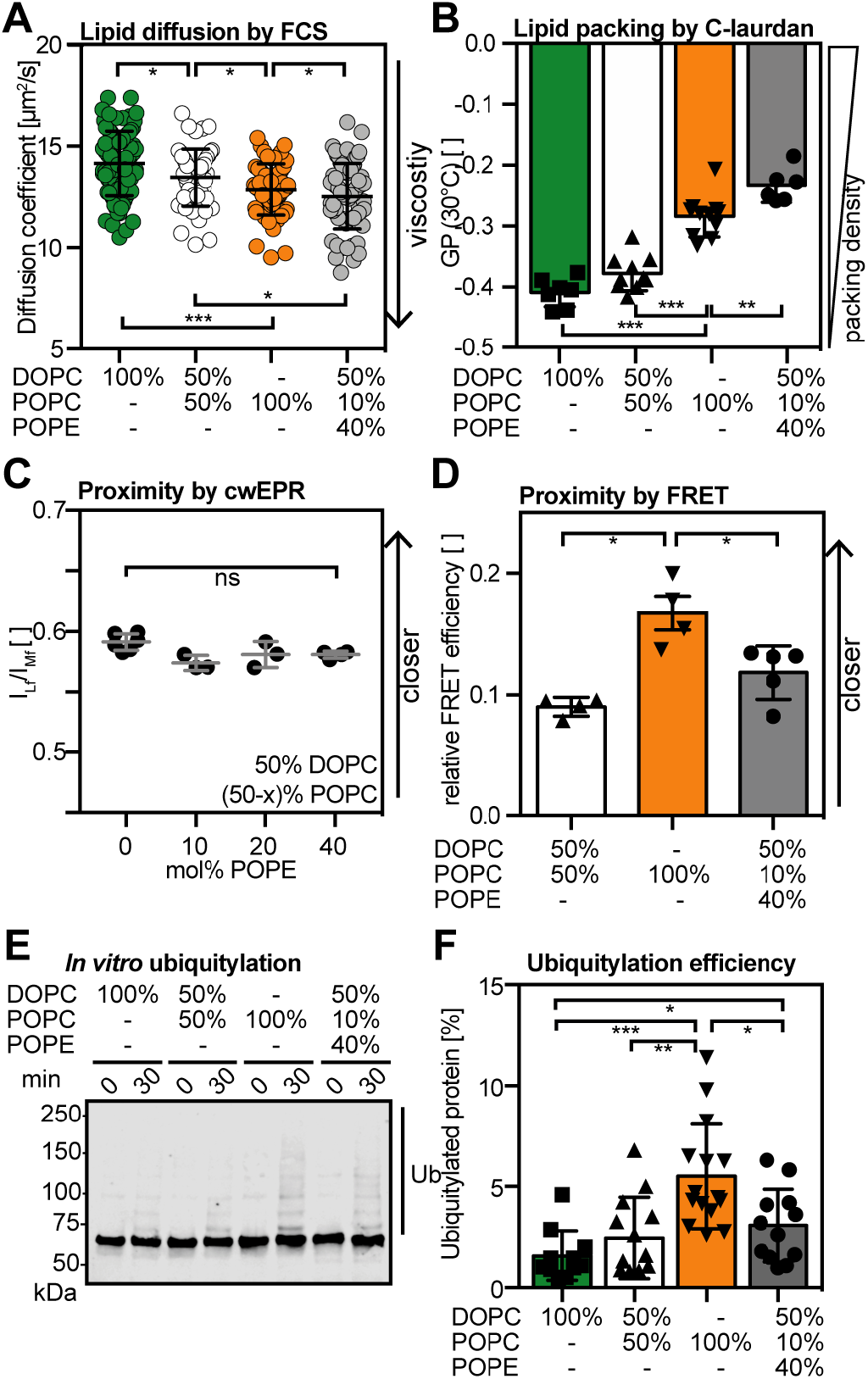
The conformation and activity of the sense-and-response construct does not correlate with membrane viscosity. **(A)** Diffusion coefficients of Atto488-DPPE in giant unilaminar vesicles of the indicated lipids were determined by confocal point FCS. The data are shown as mean ± SD (n ≥ 52). A Kolmogorov-Smirnov test was performed to test for statistical significance (*p<0.05, ***p<0.001). **(B)** The lipid packing of liposomes with indicated lipid compositions was determined by C-laurdan spectroscopy at 30°C. The index of lipid packing is represented as generalized polarization (GPs) ranging from +1 for most ordered to −1 for most disordered membrane lipids. The data are shown as mean ± SD (n ≥4 as indicated) A two-tailed unpaired t-test was performed to test for statistical significance (**p<0.01, ***p<0.001). **(C)** cwEPR spectra were recorded at −115°C for a fusion protein composed of MBP and the TMH of Mga2 (^MBP^Mga2^1032-1062^) labeled at position W1042C and reconstituted at a molar protein:lipid of 1:500 in liposomes composed of the indicated lipids. The semi-quantitative proximity index I_Lf_/I_Mf_ was derived from the cwEPR spectra as in ^26^. Higher values indicate a lower average interspin distance. Plotted is the mean ± SD (n ≥ 3 as indicated). A two-tailed unpaired t-test was performed to test for statistical significance (^ns^p≥0.05). **(D)** Relative FRET efficiencies calculated from fluorescence emission spectra (ex: 488 nm, em: 500-700 nm) of the (K983^D^+K969^A^) FRET pair reconstituted in liposomes composed of 50 mol% DOPC, 10 mol% POPC and 40 mol% POPE. The relative FRET efficiencies measured in 50 mol% DOPC, 50 mol% POPC and 100 mol% POPC (same as in Figure 2G) are shown for comparison. Shown are mean ± SD (n ≥ 4 as indicated). A two-tailed unpaired t-test was performed to test for statistical significance (*p<0.05). **(E)** *In vitro* ubiquitylation of the zipped sense-and-response construct (^ZIP-MBP^Mga2^950-1062^) reconstituted in liposomes of the indicated lipid compositions at a molar protein-to-lipid ratio of 1:8,000. After the reaction was stopped, the samples were subjected to SDS-PAGE and analyzed by immunoblotting using anti-MBP antibodies. **(F)** Densiometric quantification of ubiquitylated species from the *in vitro* ubiquitylation assay at the indicated time points from immunoblots as in (E). Plotted is the mean ± SD (n ≥11 as indicated). A two-tailed unpaired t-test was performed to test for statistical significance (*p<0.05, **p<0.01, ***p<0.001).

If Mga2 would directly sense membrane viscosity, the fluidity of the bilayer should dominate the structural dynamics of the sensory TMHs and at the site of ubiquitylation (Figure 3). Most importantly, the membrane viscosity should then also control the ubiquitylation of the sense-and-response construct (Figure 2). Using up to 40 mol% of PE in the lipid bilayer to perturb the membrane viscosity without changing the composition of its lipid acyl chains, we rigorously tested these predictions (Figure 4). (i) We found no evidence that different proportions of PE in the bilayer perturb the conformational dynamics in the sensory TMH of Mga2, when studied by EPR spectroscopy (Figure 4C, S3A). We took advantage of a previously established minimal sensor construct, which comprises the TMH of Mga2 (residues 1029-1062) fused to MBP^26^. Using methanethiosulfonate (MTS) spin labels installed at the position of W1042 in the TMH and continuous wave EPR spectroscopy, we had previously observed a significant impact of lipid saturation on the observed interspin distances^26^. Here, we show that up to 40 mol% of PE in the bilayer has no discernable impact on the resulting EPR spectra (Figure S3B) and the semiquantitative value for average interspin proximity (the ILf/IMf ratio) (Figure 4C) even though it significantly increases membrane viscosity (Figure 4A). This means that the previously reported impact of lipid saturation on the structural dynamics of the TMH^26^ cannot be caused by increased membrane viscosity (Figure 4A). (ii) The role of membrane viscosity on the structural dynamics in the region of Mga2 ubiquitylation was addressed using our newly established FRET reporter (K983^D^+K969^A^) (Figure 3). The FRET efficiency reports on the average proximity between the binding site of the E3 ubiquitin ligase Rsp5 (K969^A^) and a target site of ubiquitylation (K983^D^) in the opposing protomer of Mga2. The FRET efficiency of the reporter placed in a bilayer with 40 mol% PE was moderately higher than in a PE-free bilayer with an otherwise identical acyl chain composition (50 mol% DOPC, 50 mol% POPC). More strikingly, the highest FRET efficiency was observed in a more saturated membrane environment (POPC), which is less viscous than the PE-containing membrane (Figure 4A, D, S3C). Thus, the FRET efficiency of this reporter does not correlate with membrane viscosity. (iii) The functional relevance of membrane viscosity was tested by studying its impact on the *in vitro* ubiquitylation of the sense-and-response construct. The highest degree of ubiquitylation was observed in a POPC bilayer, which also has the highest degree of lipid saturation (Figure 4E, F). For a PE-containing bilayer, which is less saturated but more viscous, we observed significantly less ubiquitylation (Figure 4E, F). Together, these structural and functional data indicate that a key mediator of the homeoviscous response in baker’s yeast does not sense membrane viscosity. Instead, they highlight a particular sensitivity of Mga2 to the degree of lipid saturation.

### The configuration and position of lipid unsaturation controls the output signal of Mga2

To gain deeper insight into how the double bond in unsaturated lipid acyl chains might contribute to the activation of Mga2, we employed a different set of lipids. We used PC lipids with two unsaturated (18:1) acyl chains differing either in the position (Δ6 or Δ9) or the configuration (Δ9-*cis* or Δ9-*trans*) of the double bond (Figure S4A). Expectedly, we find that the ‘kink’ introduced by a *cis* double bond supports membrane fluidity (Figure 5A, S4B) by lowering both, lipid packing and membrane order (Figure 5B). Importantly, Δ6-*cis* acyl chains render the membrane more viscous than Δ9-*cis* acyl chains (Figure 5A) with no detectable impact on membrane order as studied by C-laurdan spectroscopy (Figure 5B). In contrast, Δ9-*trans* 18:1 acyl chains render the bilayer substantially more viscous (Figure 5A) and allow for a much tighter packing of lipids (Figure 5B). Using these bilayer systems differing by the position and configuration of the double bond in the unsaturated lipid acyl chains, we set out to study their impact on various aspects of the structure and function of Mga2 *in vitro*.

**Figure 5:**
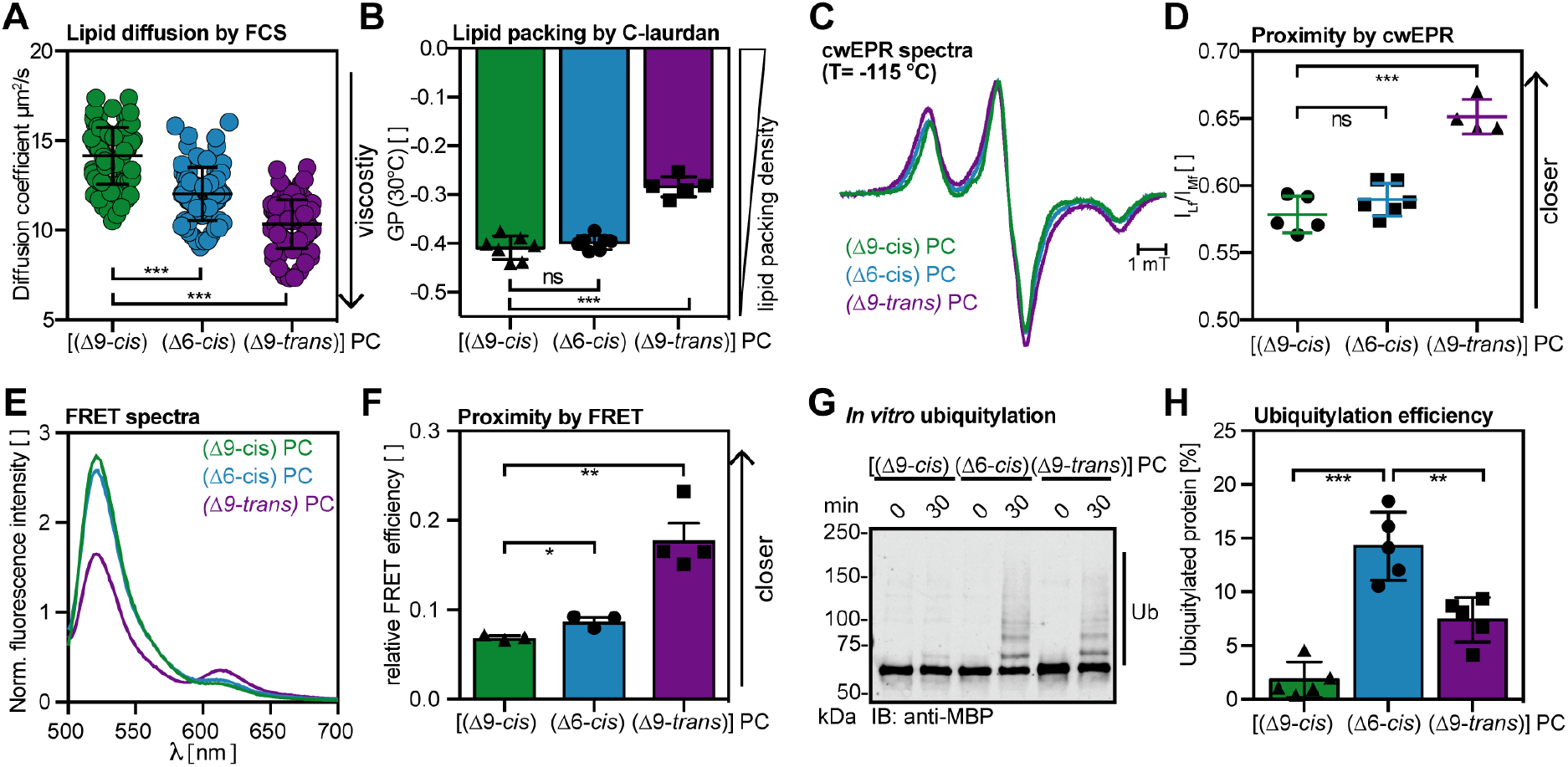
The position and configuration of the double bond in the lipid acyl chains affects the configuration and activity of the sense-and-response construct. **(A)** Diffusion coefficients of Atto488-DPPE in giant unilaminar vesicles of the indicated lipids were determined by confocal point FCS. The data are shown as mean ± SD (n ≥ 84). A Kolmogorov-Smirnov test was performed to test for statistical significance (***p<0.001). **(B)** The lipid packing in liposomes with the indicated lipid compositions was determined by C-laurdan spectroscopy at 30°C. The index of lipid packing is represented as generalized polarization (GPs) ranging from +1 for most ordered to −1 for most disordered membrane lipids. The data are shown as mean ± SD (n ≥4 as indicated). An unpaired two-tailed students t-test was performed to test for statistical significance (***p<0.001) **(C)** Intensity normalized cwEPR spectra recorded at −115°C for a fusion protein composed of MBP and the TMH of Mga2 (^MBP^Mga2^1032-1062^) labeled at position W1042C was reconstituted at a molar protein:lipid of 1:500 in liposomes composed of the indicated PC lipids. **(D)** The semi-quantitative proximity index ILf/IMf was derived from cwEPR spectra as in ^26^. Higher values indicate a lower average interspin distance. Plotted is the mean ± SD (n ≥ 4 as indicated). A two-tailed unpaired t-test was performed to test for statistical significance (***p<0.001). **(E)** Fluorescence emission spectra of the (K983^D^+K969^A^) FRET pair reconstituted in liposomes composed of different PC lipids were recorded (ex: 488 nm, em: 500-700 nm), normalized to the maximal acceptor emission after direct acceptor excitation (ex: 590 nm), and plotted. The emission spectra were normalized to acceptor emission after direct acceptor excitation. The emission spectrum measured in DOPC is shown for comparison (same as in Figure 2F). **(F)** Relative FRET efficiencies calculated from spectra as in (E). Shown are mean ± SD (n ≥3 as indicated). A two-tailed unpaired t-test was performed to test for statistical significance (*p<0.05, **p<0.005). The relative FRET efficiencies measured in 100 mol% DOPC (same as in Figure 2G) is shown for comparison. **(G)** *In vitro* ubiquitylation of the zipped sense-and-response construct (^ZIP-MBP^Mga2^950-1062^) reconstituted in liposomes of the indicated lipid compositions at a molar protein-to-lipid ratio of 1:8,000. After the reaction was stopped, the samples were subjected to SDS-PAGE and analyzed by immunoblotting using anti-MBP antibodies. **(H)** Densiometric quantification of ubiquitylated species from the *in vitro* ubiquitylation assay at the indicated time points from immunoblots as in (G). Plotted is the mean ± SD from five independent experiments.

First, we studied how the double bond position and configuration affects the structural dynamics of Mga2’s TMH region using EPR spectroscopy (Figure 5C). A substantial broadening of the continuous wave EPR spectra recorded at −115°C (Figure 5C) and an increased interspin proximity (Figure 5D) were observed, when the sensor was placed in the tightly packed membrane with Δ9-*trans* acyl chains. Much less spectral broadening was observed in membrane environments with either Δ6-*cis* or Δ9-*cis* acyl chains (Figure 5C). This indicates that Δ9-*trans* double bonds in lipid acyl chains –more than Δ9-*cis* and Δ6-*cis* bonds – stabilize a rotational orientation of Mga2’s TMH region, where spin labels at the position W1042 face each other in the dimer interface^26^. Moreover, our data also suggests that lipid acyl chains with Δ9-*trans* double bonds, which are less kinked than those with Δ9-*cis* double bonds, have a similar impact on the structural dynamics of Mga2’s TMH as saturated lipid acyl chains.

Next, we tested if the position and configuration of the double bond in unsaturated lipid acyl chains has an impact on the structural dynamics of Mga2 in the region of ubiquitylation. To this end, we used our newly established FRET reporter (K983^D^+K969^A^) and reconstituted it successfully in these new lipid compositions (Figure S4C). The FRET signal (Figure 5E) and FRET efficiency (Figure 5F) of the reporter was low when it was situated in a bilayer with poorly packing Δ9-*cis* acyl chains (Figure 2F, G and Figure 5E, F). This indicates a relatively large distance between the binding site for Rsp5 (K969) and target site for ubiquitylation in opposing protomer of Mga2 (K983). The FRET efficiency was slightly higher, when the reporter was reconstituted in a membrane composed of lipids with Δ6-*cis* acyl chains (Figure 5F). This suggests that the position of the *cis*-double bond has a significant, but rather modest impact on the average distance between K969^A^ and K983^D^ in the FRET reporter. The highest FRET efficiency was observed when the reporter was placed in a membrane with tightly packing Δ9-*trans* 18:1 acyl chains (Figure 5F). These findings demonstrate that the structural dynamics of Mga2 is affected by the position and configuration of the double bond in unsaturated lipids. Furthermore, our data suggest a robust, structural coupling between the TMH of Mga2 (Figure 5C, D) and site of ubiquitylation (Figure 5E, F).

In order to address the functional consequences of these structural changes, we performed *in vitro* ubiquitylation assays with the sense-and-response construct (^ZIP-MBP^Mga2^950-1062^) reconstituted in the three distinct membrane environments. While barely any ubiquitylation above background was detectable, when the sense-and-response construct was reconstituted in a loosely packed bilayer with Δ9-*cis* acyl chains, we observed a robust ubiquitylation when the construct was situated in a bilayer with either Δ9-*trans* or Δ6-*cis* acyl chains. Strikingly, the highest degree of ubiquitylation of the reporter was observed in the membrane with Δ6-*cis* lipid acyl chains, followed by the more viscous and more tightly packed membrane with Δ9-*trans* lipid acyl chains. This observation supports our previous conclusion that the ubiquitylation of Mga2 does not correlate with membrane viscosity. Furthermore, these data establish that Mga2 does not sense the mere presence or absence of double bonds in the lipid acyl chains. Instead, it is highly sensitive to the configuration and position of the double bond with its immediate effect on the structural and dynamic properties of the lipid acyl chains, which ultimately seem to dictate the ubiquitylation of Mga2.

### The bulkiness of the sensory residue in Mga2 determines the signal output

The TMH of Mga2 contains a bulky tryptophan (W1042), which is functionally important and might serve as sensor residue^26^. Previous MD simulations have shown that this residue is situated in the hydrophobic core of the bilayer overlapping with the Δ9-*cis* double bonds of unsaturated phospholipids^26^. We hypothesize that Mga2 might sense a thin slice of the lateral pressure or lateral compressibility profile^2,20^. In fact, the sensitivity of our sense-and-response construct to the position of the double bond in unsaturated lipids (Figure 5) is consistent with this idea. Our model predicts that the activation of Mga2 is controlled by the size of the amino acid side chain at position 1042, which also controls the population of alternative, rotational configurations of the sensory TMH in a dynamic equilibrium. An increased lateral pressure/compressibility (e.g. by increased lipid saturation) in the region should cause sizeable amino acids to ‘hide’ in the dimer interface thereby stabilizing a productive configuration. A smaller residue should be less sensitive to the membrane environment and populate non-productive configurations.

In order to test this prediction, we have substituted W1042 to either tyrosine (Y), phenylalanine (F), glutamine (Q), leucine (L), or alanine (A) and assayed the role of the sidechain bulkiness and aromatic character on the signal output *in vivo*. Expectedly, a Δ*SPT23ΔMGA2* double mutant lacking both transcriptional regulators of *OLE1* does not grow unless unsaturated fatty acids (UFAs) were provided with the medium (Figure 6A)^48^. This UFA auxotrophy of Δ*SPT23ΔMGA2* cells is complemented by both wild type and mutant *MGA2* variants expressed from the endogenous promotor on a *CEN*-based plasmid (Figure 6A, B). However, the growth of these cells was highly dependent on the amino acid at position 1042 under UFA-limiting conditions (Figure 6A). Furthermore, we observed a striking correlation between the size of the side chain and the optical density of overnight cultures (Figure 6B). The only exception to this near-perfect correlation was the W1042Q mutation. Given that intra-membrane glutamines are known to mediate homotypic interactions^49^, we speculate that the W1042Q mutation stabilizes a rotational conformation of the TMHs, where the two Q1042 side chains face each other and interact, thereby stabilizing Mga2 in a processing-competed configuration. Intriguingly, the phenotypic differences between the W1042Q, W1042L and W1042A variants show that an aromatic character at the sensory position is not absolutely required for sensing.

**Figure 6:**
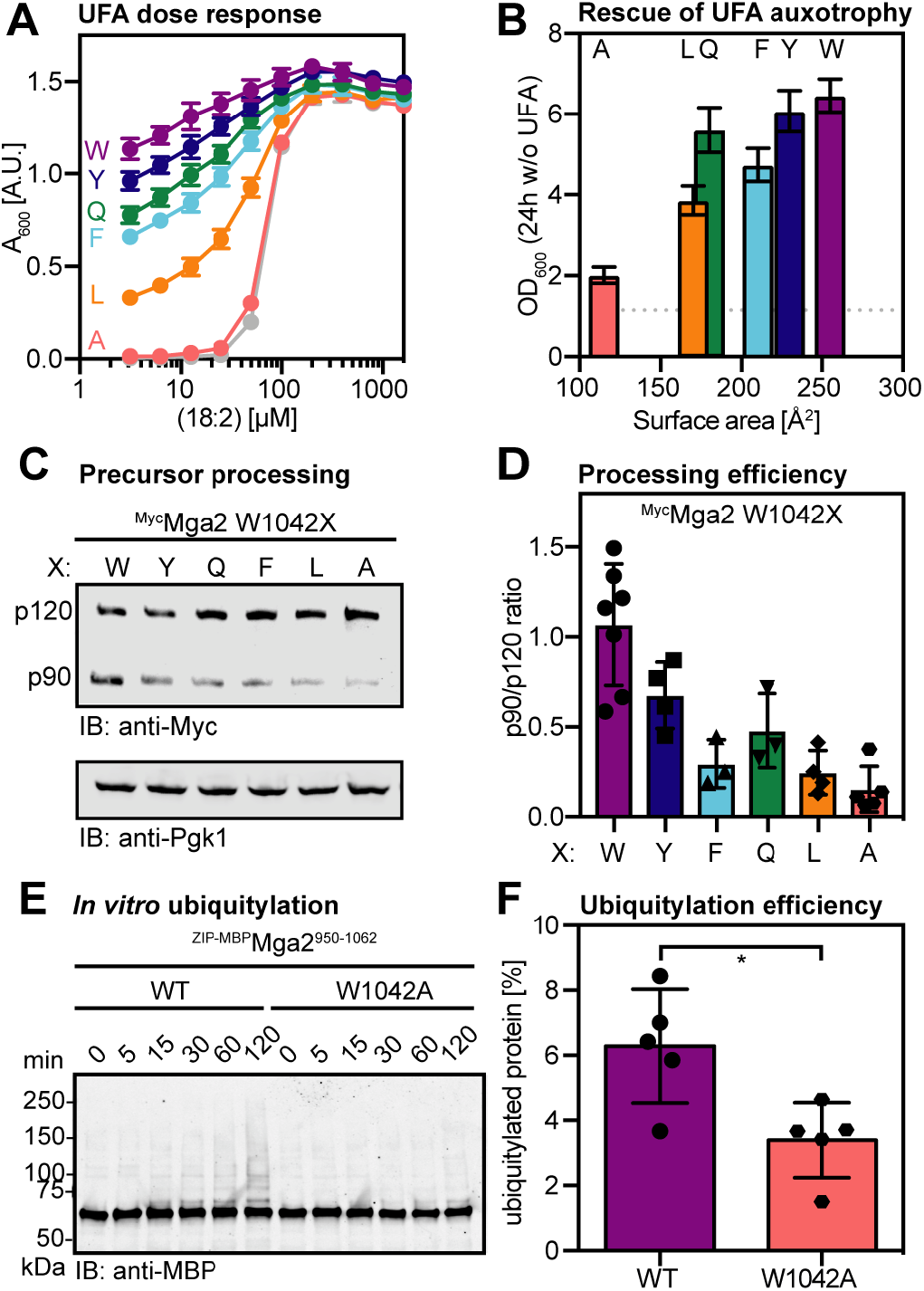
The size and polarity of the TMH residue 1042 affects the activity of Mga2 *in vivo* and *in vitro*. **(A)** Dose-dependent rescue of UFA auxotrophy by linoleic acid (18:2). *ΔSPT23ΔMGA2* strains carrying *CEN*-based plasmids to produce ^Myc^Mga2 variants with the indicated residues at position 1042 were cultivated for 16 h at 30°C in SCD-Ura medium supplemented with indicated concentrations of linoleic acid in 0.8% tergitol. The density of the culture was determined at 600 nm (OD_600_) and plotted against the concentration of linoleic acid. Cells carrying an empty vector served as control (gray). Plotted is the mean ± SEM (n = 8). **(B)** Rescue of UFA auxotrophy of Δ*SPT23*Δ*MGA2* by Mga2 variants. Cells producing mutant Mga2 as in (A) were cultivated for 24 h in the absence of supplemented UFAs in SCD-Ura medium. Cell density was determined as in (A) and plotted against residue surface area of residues installed at position 1042 ^62^. Plotted is the mean ± SEM of five independent experiments. The dotted line indicates the OD measured for an empty vector control. **(C)** Immunoblot analysis of the Mga2 processing efficiency. Wild type cells (BY4741) producing the indicated ^Myc^Mga2 variants at position 1042 were cultivated in YPD to the mid-exponential phase. Cell lysates were subjected to SDS-PAGE and analyzed via immunoblotting using anti-Myc antibodies to detect the unprocessed (p120) and the processed, active form (p90) of Mga2. An immunoblot using anti-Pgk1 antibodies served as loading control. **(D)** Densiometric quantification of the ratio of p90:p120 in immunoblots as in (C). Plotted is the mean ± SD (n ≥ 3 as indicated). **(E)** *In vitro* ubiquitylation of the zipped sense-and-response construct ^ZIP-MBP^Mga2^950-1062^ wild type (WT) and a W1042A variant reconstituted at a protein:lipid molar ratio of 1:15,000 in POPC. After the reaction was stopped, ubiquitylated species were detected by SDS-PAGE and subsequent immunoblotting using anti-MBP antibodies. (F) Densiometric quantification of the *in vitro* ubiquitylation assays as in (E). The fraction of ubiquitylated protein was determined for the indicated time points and for the wildtype (WT) and W1042A variant of the sense-and-response construct. Plotted is the mean ± SD (n = 5). The statistical significance was tested by a two-tailed, unpaired t-test (*p<0.05).

Next, we studied the impact of these mutations on the proteolytic processing of full-length Mga2 in cells (Figure 6C, D). We found a perfect concordance of these immunoblot experiments with the *in vivo* phenotypes (Figure 6A, B). The processing of the membranebound precursor of Mga2 (P120) to the signaling-active form (P90) was greatly affected by the residue at the position 1042. These data were complemented by functional *in vitro* experiments using the sense-and-response construct (Figure 6E, F). The *in vitro* ubiquitylation of the sense-and-response construct reconstituted in a POPC bilayer was significantly reduced to almost background levels by the W1042A mutation. Based on these *in vivo* and *in vitro* data, we conclude that the size and the chemical character of the amino acid at position 1042 is of central importance for the signaling output.

## Discussion

We have reconstituted key steps of sensing and communicating lipid saturation by the prototypical type II membrane property sensor Mga2^21^. We uncover a unique sensitivity Mga2 to the lipid acyl chain composition of the ER membrane and provide direct evidence for a functional coupling between the dimeric, sensory TMHs and the sites of ubiquitylation. Our *in vitro* system allowed us to directly test a central assumption underlying the concept of homeoviscous adaptation^9,15,17^. By investigating the role of membrane viscosity on the ubiquitylation of Mga2, we demonstrate that the core regulator of fatty acid desaturation in baker’s yeast^23,50^ is not regulated by membrane fluidity (Figure 4, 5). Instead, our data suggest that Mga2 uses a bulky TMH residue (W1042) to sense a thin slice of the lateral pressure/compressibility profile in a specific region of ER membrane. Based on our findings, we conclude that membrane fluidity does not serve as the central measured variable for regulating the lipid acyl chain composition in baker’s yeast and presumably many other eukaryotic species.

Our *in vitro* approach with reconstituted proteoliposomes has provided unprecedented insights into the sensitivity of Mga2 to physiologically relevant changes of the lipid acyl chain composition^8,44^. The sense-and-response construct cannot be ubiquitylated in a relatively unsaturated membrane (75% Δ9-*cis* 18:1 acyl chains), but it is robustly ubiquitylated in a slightly more saturated environment (50% Δ9-*cis* 18:1 acyl chains) (Figure 2F). A simple back-of-an-envelope calculation that considers only the volume of the lipid bilayer highlights the remarkable dose-response relationship of this machinery: The sense-and-response system is *OFF*, when the concentration of unsaturated lipid acyl chains is ~1.9 M, but it is *ON* at a concentration of ~1.3 M (assumptions: ~370,000 lipids per 200 nm liposome; ~4,82×10^−19^ l membrane volume per liposome). This switch-like response is based on fluctuating signals from the membrane, which are decoded by the sensor protein into an almost binary output.

Our results lead to the following model of lipid saturation sensing: The lipid acyl chain composition has profound impact on the lateral pressure/compressibility profile^51^, which determines the population of alternative rotational orientations in the transmembrane helix region of Mga2, as previously proposed^26^ and supported by our EPR data (Figure 4C,D). In a more saturated membrane, the sensory tryptophan (W1042) points more likely towards the dimer interface, while in a more unsaturated membrane it points more often away from the interface towards the lipid environment^26^. Nevertheless, in any fluid bilayer the dimeric TMHs constantly rotate against each other and explore various alternative rotational states. The fluctuating signal from the membrane is thus encoded by the structural dynamics of the TMHs, which is then transmitted to the sites of ubiquitylation and E3 ubiquitin ligase binding via a disordered region (Figure 3F,G). We speculate that the flexible linkage provides a means to bias the orientation and relative position of two ‘ubiquitylation zones’ around the E3 ubiquitin ligase Rsp5 bound to Mga2, however, with a minimal perturbation of the TMH-dynamics. Such ‘zones of ubiquitylation’ have recently been predicted for Rsp5 and implicated into the quality control of misfolded and mistargeted plasma membrane proteins^52^. Supported by our FRET data (Figure 4), we propose that Rsp5 bound to one protomer of dimeric Mga2 can ubiquitylate specific lysine residues on the other, when it is properly placed and oriented. This *trans*-ubiquitylation would effectively be controlled by the physicochemical properties of the ER membrane. The remarkable sensitivity of Mga2 ubiquitylation to the lipid environment might be sharpened by deubiquitylating enzymes^54^ such as Ubp2^53^ and supported by an activating, *trans*-autoubiquitylation of the Rsp5^55^.

The assays and tools established here, provide new handles to better understand the structural and dynamic features that render a protein a good substrate of the E3 ubiquitin ligase Rsp5. Identifying the molecular rules of substrate selection is a major open question, because Rsp5 has been implicated in most diverse aspects of cellular physiology including endocytosis^52^, mitochondrial fusion^58^, and the turnover of heat-damaged proteins in the cytosol^56^. Our *in vitro* system using a membrane-reconstituted, conditional substrate of Rsp5 provides a unique opportunity to better understand i) the contribution of *trans*-autoubiquitylation of Rsp5, ii) the relevance of structural malleability in Rsp5 substrates, and iii) the role of deubiquitylating enzymes in defining the selectivity and sensitivity of the Rsp5-mediated ubiquitylation. In the context of the Mga2 sensor, it will be most intriguing to understand how ‘noisy’ signals from the TMH region are transduced into robust, almost switch-like ubiquitylation responses.

Two lines of evidence suggest that the rotation-based sensing mechanism of Mga2^26^ is based on a collective, physical property of the membrane rather than on a preferential, chemical interaction with the double bonds in the lipid acyl chains. Firstly, Mga2 distinguishes robustly between two membrane environments that differ in the configuration of the double bonds (*cis* or *trans*) in the lipid acyl chains, but not in the overall abundance of double bonds (Figure 5). Secondly, an aromatic amino acid, which might confer some chemical specificity for double bonds, is not absolutely required at the position of the sensory tryptophan (W1042) in the TMH (Figure 6). A partial activity of the OLE pathway is preserved when the sensor residue W1042 is substituted with leucine, but not when it is substituted with the smaller alanine (Figure 6, S5). Nevertheless, our data do not rule out a contribution of chemical specificity to the sensor function. In fact, we expect that the high degree of structural malleability in the TMH region and at the site of ubiquitylation is established by a fine balance of chemical interactions and collective, physical membrane properties.

In conclusion, we have provided deep mechanistic insight into a sensory system that is centrally important for membrane adaptivity. Our findings challenge the common view of membrane viscosity as pivotal measured variable in eukaryotic cells and have important implications to all processes involving membrane lipid adaptation. Beyond that, our work represents an important step towards identifying the molecular rules of substrate selection by the E3 ubiquitin ligase Rsp5. Furthermore, this work opens a door towards establishing genetically encoded machineries that can sense specific membrane features, which are indiscernible by conventional tools. In the future, these sensors will be exploited to dissect the physical membrane properties of different organelles and cells *in vivo* and in real-time.

## Materials and Methods

All plasmids and strains used in this study are listed in Table S1 and S2. For detailed description of experimental procedures see Supplementary Materials.

### Expression, purification and labeling of ^MBP^Mga2-fusions

The ^ZIP-MBP^Mga2^950-1062^ fusion protein comprising the leucine-zipper of the GCN4 transcription factor (residues 249-281), the MBP from *Escherichia coli*, and the residues G950–D1062 from Mga2 was generated using the pMal-C2x plasmid system. The resulting constructs were produced in *E. coli* and isolated in detergent solution using amylose affinity followed by a preparative SEC (Superdex 200 10/300 Increase). For fluorescent labeling, the K983C and K969C variants were incubated with 1 mM ATTO488 or ATTO590 (ATTO-TEC GmbH) on the affinity purification column for 16 h at 4 °C. The ^MBP^Mga2^1032-1062^ fusion protein containing residue R1032-D1062 from Mga2 and a W1042C mutation was purified and labeled with MTS spin probes as described previously ^26^. The proteins were stored in 40 mM HEPES (pH 7.0), 120 mM NaCl, 0.8 mM EDTA, 40 mM OG, and 20% (w/v) glycerol.

### Reconstitution of ^MBP^Mga2-fusions in proteoliposomes

The spin-labeled ^MBP^Mga2-TMH fusion was reconstituted at a protein:lipid molar ratio of 1:500 as described previously ^26^. The unlabeled or ATTO-labeled ^ZIP-MBP^Mga2^950-1062^ constructs were reconstituted at different protein-lipid molar ratios of 1 to 5,000, 1 to 8,000, and 1 to 15,000. To this end, lipids (final concentration 1 mM) and Octyl-β-D-glucopyranoside (final concentration 37.5 mM) were mixed with either labeled or unlabeleld proteins in a final volume of 1 ml. After 10 min of incubation at room temperature under constant agitation, the detergent was via SM-2 biobeads (two-step removal using 500 mg and 100 mg, respectively). A detailed description is provided in the SI Materials and Methods.

### Diffusion coefficients by fluorescence correlation spectroscopy

FCS on the GUVs was carried out using Zeiss LSM 880 microscope, 40X water immersion objective (numerical aperture 1.2) as described previously^59^. First, GUVs were labelled by adding fluorescent analogues to a final concentration of 10-50 ng/mL (≈0.01 mol%). To measure the diffusion on the GUV membrane, vesicles were placed into an 8-well glass bottom (#1.5) ibidi chambers coated with BSA. GUVs of small sizes (≈10 *μ*m) were picked for measurements. The laser spot was focused on the top membrane of the vesicles by maximizing the fluorescence intensity. Then, 3-5 curves were obtained for each spot (five seconds each). The obtained curves were fit using the freely available FoCuS-point software using 2D and triplet model ^60^.

### C-laurdan spectroscopy

C-laurdan was used to measure lipid packing^47^. To this end, 333.3 *μ*M lipid was mixed with 0.4 *μ*M C-laurdan dye in 150 *μ*l 50 mM HEPES pH 7.4, 150 mM NaCl, 5 %(w/V) glycerol. The sample was excited at 375 nm and an emission spectrum from 400 to 600 nm was recorded (excitation and emission bandwidth 3 nm). For blank-correction, an emission spectrum recorded in the absence of C-laurdan was used. The generalized polarization (GP) value was calculated by integrating the intensities between 400 – 460 nm (I_Ch1_) and 470 – 530 nm (I_Ch2_).

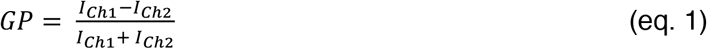

### Recording and analysis of FRET spectra

For FRET measurements, the ZIP-MBPMga2950-1062 K983ATTO488 and ZIP-MBPMga2950-1062 K969^ATTO590^ constructs were used as fluorescence donor and acceptor, respectively. Fluorescence emission spectra were recorded in detergent solution and in proteoliposomes at 30°C. The samples were excited at 488 nm and 590 nm for donor and acceptor excitation, respectively. The spectra were normalized to the maximal acceptor fluorescence intensity after direct excitation to correct for subtle variations in the reconstitution yields. Since the bleed-through for both, donor and acceptor fluorescence was negligible, ratiometric FRET (relative FRET: Erel) was determined as the donor-to-acceptor intensity ratio at 525 nm and 614 nm from the raw data (equation 2) for qualitative comparisons.

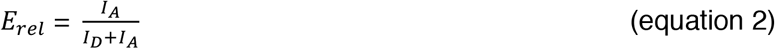

### *In vitro* ubiquitylation assay

Proteoliposomes containing ^ZIP-MBP^Mga2^950-1062^, ^8xHis^Ubiquitin (see SI Materials and Methods for a description of expression and purification), cytosol, and an 10x ATP regenerating system were mixed on ice in a total volume of 20 *μ*l to obtain final concentrations of 0.1 *μ*M ^ZIP-MBP^Mga2^950-1062^, 0.1 *μ*g/*μ*l ^8xHis^Ubiquitin, 1 *μ*g/*μ*l cytosolic proteins, 1 mM ATP, 50 mM creatine phosphate and 0.2 mg/ml creatine phosphokinase in ubiquitylation buffer (20 mM HEPES, pH 7.4, 145 mM NaCl, 5 mM MgCl_2_, 10 *μ*g/ml chymostatin, 10 *μ*g/ml antipain, 10 *μ*g/ml pepstatin). Cytosol was prepared from BY4741 cells grown to mid-log phase (OD_600_ = 1) in YPD medium as previously described^42^. The ubiquitylation reaction was incubated at 30 °C and stopped by mixing the sample at a ratio of 2:1 with 5x reducing sample buffer (8 M urea, 0.1 M Tris-HCl pH 6.8, 5 mM EDTA, 3.2% (w/v) SDS, 0.15% (w/v) bromphenol blue, 4% (v/v) glycerol, 4% (v/v) β-mercaptoethanol) and boiling it. Protein ubiquitylation was analyzed by SDS-PAGE using 4-15% Mini-PROTEAN-TGX gels (BioRad) and immunoblotting using anti-MBP antibodies.

## Acknowledgments

We thank Laura Glück and Kim Wendrich for excellent technical assistance, and Roberto Covino and Ilya Levental for critically reading the manuscript. We like to acknowledge Jeffrey Brodsky and Volker Dötsch for sharing reagents and protocols as well as Roberto Covino and Gerhard Hummer for ongoing fruitful discussions. This work was supported by the Deutsche Forschungsgemeinschaft (DFG, ER608/2-1) to R.E. and (HA6322/3-1) to I.H., the Volkswagen Foundation (Life?, grant no. 93089) to R.E., and the European Molecular Biology Organization (EMBO, ASTF 451-2014) to S.B.. ES is funded by British Council Newton-Katip Celebi Fund (#352333122).

## Author contributions

Conceptualization: R.E.; Experimental Design: S.B., E.S., D.W., I.H., R.E.; Performed experiments: S.B., E.S., D.W.; Writing – original draft: S.B., R.E.; Funding Acquisition: R.E., I.H., E.S.; Supervision: R.E..

## Data and materials availability

All data needed to evaluate the conclusions in the paper are present in the paper and/or Supplementary Materials. Additional data and materials related to this paper may be requested from the authors.

## Supplementary Materials

Fig. S1. Isolation and functional reconstitution of sense-and-response construct.

Fig. S2. Establishing a FRET reporter based on sense-and-respond construct.

Fig. S3. Reconstituting sense-and-response construct in PE-containing liposomes.

Fig. S4. Reconstituting sense-and-response construct in liposomes with different PC-species.

Fig. S5. Mutagenesis of sensory residue W1042 and phenotypic characterization.

Table S1. Plasmids used in this study.

Table S2. Strains used in this study.

### Reagents and antibodies

All chemicals and reagents were of analytical or higher grade and obtained from Sigma Aldrich if not stated otherwise. The following antibodies were used: mouse anti-Myc (9E10), mouse anti-Pgk1 (Life Technologies), mouse anti-MBP (NEB), anti-mouse-HRP (Dianova), anti-mouse-IRDye 800CW (LI-COR). Atto488-PE was purchased from AttoTec GmbH. Abberior Star Red-Cholesterol is purchased from Abberior GmbH. It has a PEG linker between cholesterol moiety and the fluorescent tag.

### Cultivation and genetic manipulation of *S. cerevisiae*

Overnight cultures were inoculated from single colonies and cultivated in SCD selection medium at 30°C until the stationary phase was reached. The UFA auxotroph *ΔSPT23ΔMGA2* strain was cultivated in the presence of 0.05% sodium linoleate. Main cultures were inoculated to an OD_600_ of 0.2 in rich medium (YPD) and cultivated to the mid-exponential phase (OD_600_ ≈ 1.0). If indicated, the YPD was supplemented with sodium linoleate.

A *CEN*-based plasmid expressing 3xmyc-tagged *MGA2* under the control of the *MGA2* promotor for near-endogenous levels was used as described previously ^26^. Mutagenesis of *MGA2* was performed using a PCR-based strategy based on the QuikChange^®^ method (Stratagene) using the PHUSION polymerase (NEB). *S. cerevisiae* was transformed using Lithium-Acetate (Ito et al., 1983).

### Preparation of cell extracts and immunoblot analysis

Crude cell lysates were prepared as described previously ^26^ with minor modifications. Shortly, 15 OD_600_ equivalents of cells grown to the mid-exponential phase (OD_600_ ≈ 1.0) were harvested by centrifugation, washed with phosphate-buffered saline (PBS) supplemented with 10 mM NEM and snap-frozen. The cells were resuspended in 0.5 ml lysis buffer (PBS, 10 mM NEM, 5 mM EDTA, 10 *μ*g/ml chymostatin, 10 *μ*g/ml antipain, 10 *μ*g/ml pepstatin) and lysed by bead-beating twice with 200 *μ*l zirconia beads (Roth) using a Scientific Industries SI^TM^ Disruptor Genie^TM^ Analog Cell Disruptor for 5 min each at 4 °C and 1 min pause on ice. For protein denaturation the extract was mixed at a ratio of 2:1 with 5x reducing sample buffer (8 M urea, 0.1 M Tris-HCl pH 6.8, 5 mM EDTA, 3.2% (w/v) SDS, 0.15% (w/v) bromphenol blue, 4% (v/v) glycerol, 4% (v/v) β-mercaptoethanol) and incubated at 60°C for 10 min. Centrifugation (1 min, 16,000x g, room temperature) cleared protein samples were subjected to a discontinuous SDS-PAGE using 4-15% Mini-PROTEAN-TGX gels (BioRad). After semi-dry Western-Blotting onto nitrocellulose membranes, the target proteins were detected using specific antibodies.

### Yeast growth assays / rescue of UFA auxotrophy

The UFA auxotroph Δ*SPT23*Δ*MGA2* strain was generated by Harald Hofbauer (Graz University) and cultivated in SCD-medium supplemented with 0.05% sodium linoleate. The cells were harvested by centrifugation, washed successively with 1% NP40-type tergitol (NP40S Sigma), then ddH2O and then resuspended in SCD medium lacking any additives to an OD_600_ of 0.2. The cells were either cultivated at 30°C for 5-6 h to starve cells for UFAs prior to perform spotting tests or for 24 h to study the impact of mutations on the final cell density in liquid culture. For spotting tests, the UFA-starved cells were harvested and adjusted to an OD_600_ of 1. Serial 1:10 dilutions were prepared (10^0^, 10^−1^,10^−2^, 10^−3^) and 5 *μ*l of each dilution were spotted onto selective agar plates. The plates were incubated for 2-3 days at 30°C until sufficient cell growth became apparent.

The impact of linoleate on the final cell density in liquid medium was tested with UFA-depleted cultures that were adjusted to an OD_600_ of 0.05. 50 *μ*l of these cultures were added to 180 *μ*l SCD-Ura containing 1% NP40-type tergitol and varying concentrations of linoleic acid. The optical density of the cultures was determined using a microplate reader at 600 nm (OD_600_) after 17 h of cultivation at 30°C.

### Expression, purification and labeling of ^MBP^Mga2-fusions

The minimal sensor construct (^MBP^Mga2^1032-1062^) comprising the residues R1032-D1062 that include the TMH region of Mga2 was described previously ^26^. The sense-and-response construct (^MBP^Mga2^950-1062^) was generated by cloning the coding regions of the JM and TMH region of Mga2 (residues 950-1062) into the pMal-C2x vector. The ^ZIP-MBP^Mga2^950-1062^ construct was generated by fusing the leucine zipper sequence derived from the *GCN4* transcription factor (residues 249-281) in frame to MBP protein. The minimal sensor construct and the sense-and-response construct were overexpressed in the cytosol of *E. coli* BL21(DE3)pLysS and isolated essentially as described previously ^26,63^ with minor modifications. A 500 ml culture in LBrich medium (LB medium supplemented with 2% glucose, 100 mg/ml ampicillin, 34 *μ*g/ml chloramphenicol) was inoculated 1:50 using an overnight culture and cultivated at 37°C until an OD_600_ of ~0.6 was reached. Then, protein production was induced by isopropyl-β-D-thiogalactopyranoside (IPTG) at a final concentration of 0.3 mM. After 3 h of cultivation at 37 °C the cells were harvested by centrifugation and washed with PBS. For isolation of the proteins, the cells were resuspended in 40 ml of lysis buffer (50 mM HEPES pH 7.0, 150 mM NaCl, 1 mM EDTA, 10 *μ*g/ml chymostatin, 10 *μ*g/ml antipain, 10 *μ*g/ml pepstatin, 2 mM DTT, 5 U/ml Benzonase) per liter of culture and disrupted by sonification using a SONOPULS HD2070 ultrasonic homogenizer (Bandelin) (4x 30s, power 30%, pulse 0.7 sec/0.3 sec). The protein was solubilized by gentle agitation in the presence of 50 mM b-Octylglucoside (β-OG) for 20 min at 4 °C. Non-solubilized material was pelleted by centrifugation (30 min, 100,000 x g, 4° C) and the supernatant was applied to washed and equilibrated amylose beads (NEB) using 6 ml of slurry per liter of culture. After binding (20 min at 4 °C) to the amylose column and washing the column with 26 column volumes (CV) wash buffer (50 mM HEPES pH 7.0, 200 mM NaCl, 1 mM EDTA, 50 mM β-OG) the protein was either labeled or directly eluted. The labeling of the proteins at single cysteine residues with 1 mM MTS (methanethiosulfonate) (Enzo Life Sciences) or 1 mM ATTO488/ATTO590 dyes (ATTO TEC GmbH) was performed on the amylose column during an overnight incubation at 4 °C including gentle shaking. This step was skipped for the isolation of unlabeled proteins. The fusion protein was eluted with elution buffer (50 mM HEPES pH 7.0, 150 mM NaCl, 1 mM EDTA, 10 mM maltose, 50 mM β-OG). The sense-and-response construct (^ZIP-MBP^Mga2^950-1062^) was further purified by preparative SEC using a Superdex 200 10/300 increase column in SEC-buffer (50 mM HEPES pH 7.0, 150 mM NaCl, 1 mM EDTA, 50 mM β-OG). The purified proteins could be stored at −80°C for extended periods of time in storage buffer (40 mM HEPES pH 7.0, 120 mM NaCl, 0.8 mM EDTA, 40 mM β-OG, and 20% (v/v) glycerol).

The efficiency of spin-labeling was determined for each construct by double-integration of the EPR resonances and a comparison to the signal of a 100 *μ*M MTS standard. The determined spin-label concentration was put into relation to the protein concentration determined by absorption spectroscopy at A280. The labeling efficiency for W1042C^MTS^ was > 95%.

The efficiency of labeling with fluorescent dyes was determined by absorption spectroscopy using the following extinction factors: 9.58*10^4^ l mol^−1^ cm^−1^ (unlabeled protein K983 or K969), 9*10^4^ l mol^−1^ cm^−1^ (ATTO 488), 1.2*10^5^ l mol^−1^ cm^−1^ (ATTO 590) and the correction factors were 0.1 for ATTO 488 and 0.44 for ATTO 590 according to the manufacturer’s specification. Maximal absorption intensities were determined at 505 nm (ATTO488) or 597 nm (ATTO590). The labeling efficiency was ~60% (K983^ATTO 488^) and ~90% (K969^ATTO 590^).

### Liposome preparation

Liposomes of defined compositions were generated by mixing 1,2-dioleoyl-*sn*-glycero-3-phosphocholine (DOPC), 1-palmitoyl-2-oleoyl-*sn*-glycero-3-phosphocholine (POPC), 2-dipetroselenoyl-sn-glycero-3-phosphocholine (18:1 (Δ6-*cis*)PC), 2-dielaidoyl-*sn*-glycero-3-phosphocholine (*trans*DOPC) or 1-palmitoyl-2-oleoyl-*sn*-glycero-3-phosphoethanolamine (POPE) from 20 mg/ml stocks, dissolved in chloroform to obtain following molar compositions: 1) 100% DOPC; 2) 50% DOPC/ 50% POPC; 3) 25% DOPC/ 75% POPC; 4) 100% POPC; 5) 100% 18:1 (Δ6-*cis*) PC; 6) 100% *trans*DOPC; 7) 50% DOPC/ 40% POPC/ 10% POPE; 8) 50% DOPC/ 30% POPC/ 20% POPE; 9) 50% DOPC/ 10% POPC/ 40% POPE. After evaporation of the organic solvent using a constant stream of nitrogen, the lipid film was dried in a desiccator under vacuum (2 – 4 mbar) for at least 1 h at room temperature. For rehydration, the lipid film was resuspended in reconstitution buffer (20 mM HEPES, pH 7.4, 150 mM NaCl, 5% (w/v) glycerol) to a final lipid concentration of 10 mM, incubated at 60 °C under rigorous shaking for 30 min at 1200 rpm, and incubated in a sonication in a water bath at 60°C for 30 min. The resulting multilamellar liposomes were used for reconstitution experiments.

### Reconstitution of ^MBP^Mga2-fusions in proteoliposomes

For reconstitution of the ^ZIP-MBP^Mga2^950-1062^ constructs at a protein:lipid molar ratio of 1:5,000 – 1:15,000, 0.1 *μ*mol lipid and 0.2 – 0.067 nmol protein were mixed in reconstitution buffer (20 mM HEPES (pH 7.4), 150 mM NaCl, and 5% (w/v) glycerol), adjusted to 37 mM β-OG in a total volume of 1 ml and incubated for 20 min at room temperature under gentle agitating. For detergent removal, 500 mg of Bio-Beads^TM^ SM-2 Adsorbent Media (BioRad) were added and the resulting mixture was incubated and gently mixed for 120 min at room temperature. The suspension was then transferred to a fresh tube containing 100 mg Bio-Beads^TM^ SM-2 Adsorbent Media and further incubated for 60 min. 0.8 ml of the proteoliposome containing suspension was mixed with 2.2 ml Harvesting buffer (20 mM HEPES, pH 7.4, 75 mM NaCl). Proteoliposomes were harvested by centrifugation (200,000x g, 4 °C, 18 h) and resuspended either in the respective assay buffer.

### Sucrose density gradient centrifugation

For validation of the reconstitution procedure, proteoliposomal preparation were subjected to a sucrose density step gradient and then centrifuged. To this end, 200 *μ*l of a proteoliposomal preparation were mixed with 400 *μ*l 60% (w/v) sucrose solution in reconstitution buffer and overlaid with different layers of distinct density. Depending on the protein-to-lipid molar ratio in the preparation of the proteoliposomes, two alternative step gradients were used: For protein-to-lipid molar ratios of 1:5,000 to 1:15,000, the proteoliposome-containing layer was overlaid with each 2.5 ml of 20%, 10%, 5% and 0% w/v sucrose in reconstitution buffer (gradient A). For proteoliposomes reconstituted at higher protein-to-lipid molar ratio (1:500), the proteoliposome layer was overlaid with each 3 ml of 30%, 20%, 10%, and 0% (w/v) sucrose in reconstitution buffer (gradient B). After centrifugation (100,000x g, 4°C, overnight) the gradient was fractionated from top to bottom in 0.85 ml (gradient A) or 1 ml (gradient B) fractions. The distribution of the MBP-containing fusion proteins in the gradient was analyzed by SDS-PAGE and subsequent immunoblotting. The lipid content of the individual fractions was estimated by adjusting each fraction to 7 *μ*M Hoechst 33342 and determination of the fluorescence intensity using a TECAN microplate reader (ex355 nm: em459, bandwidth 20 nm).

### Recording and analysis of cwEPR spectra

cwEPR spectra were recorded and analyzed as previously described ^26^.

### Isolation of ^His^ubiquitin

^8xHis^ubiquitin was overproduced in *E. coli* BL21(DE3)pLysS and purified using immobilized metal affinity chromatography (Ni^2+^-NTA matrix). The plasmid encoding the human ubiquitin with an N-terminal 8xHis-tag was derived from a pETM-m60 plasmid and kindly provided by the Volker Dötsch lab. The production of ^8xHis^ubiquitin was induced at an OD_600_ of ~0.6 at 37 °C using 0.3 mM IPTG. After induction, the cells were cultivated for additional 3 h at 30 °C prior to harvesting and washing of the cell pellet using PBS.

For purification, the cells were resuspended in 20 ml lysis buffer (50 mM HEPES, pH 8.0, 250 mM NaCl, 20 mM imidazol, 10 *μ*g/ml chymostatin, 10 *μ*g/ml antipain, 10 *μ*g/ml pepstatin) and disrupted by sonification (3x 30s, power 30%, pulse 0.7 s/ 0.3 s). Unbroken cells, debris, and cellular membranes were removed by centrifugation (1 h, 100,000x g, 4 °C). The cleared lysate was applied to 1 ml Ni^2+^-NTA agarose matrix and incubated for 1 h at 4 °C while rotating to allow for protein binding. The mixture was then transferred into a gravity flow column and the flow-through was collected. The affinity matrix was washed with 30 CV of wash buffer (50 mM HEPES pH 8.0, 250 mM NaCl, 20 mM imidazole). ^8xHis^ubiquitin was eluted with elution buffer (50 mM HEPES pH 8.0, 250 mM NaCl, 400 mM imidazole). The eluate was dialysed against 100-fold volume storage buffer (50 mM HEPES, pH 7.4, 150 mM NaCl) using a dialysis membrane with a molecular weight cutoff of 3.5 kDa (Spectra/Por). After 2 h the storage buffer was refreshed, and the sample was dialyzed overnight at 4°C. For long-term storage, the purified ^8xHis^ubiquitin was adjusted to 1 mg/ml and 20% (w/v) glycerol in storage buffer.

**Figure S1.**
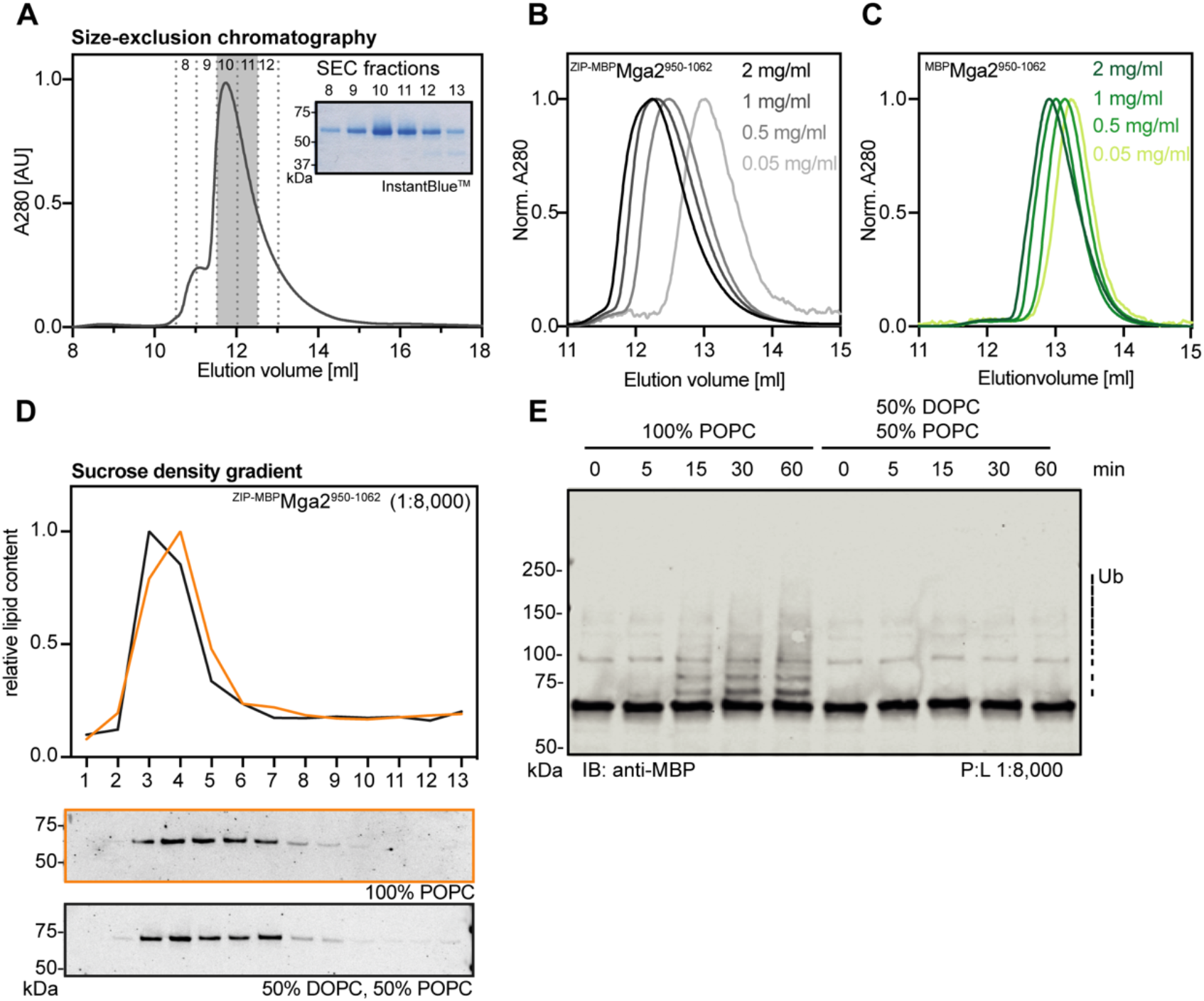
Isolation and functional reconstitution of sense-and-response construct. **(A)** Purification of the zipped sense-and-response construct (^ZIP-MBP^Mga2^950-1062^) by SEC. The eluate of the affinity purification (Figure 2C) was concentrated ~10fold and loaded onto a Superdex 200 10/300 Increase column (void volume 8.8 ml) using a 500 *μ*l loop. Fractions of 0.5 ml were collected, mixed with non-reducing membrane sample buffer and subjected to SDS-PAGE followed by InstantBlue^TM^ staining. Fraction 10 and 11 were pooled and further used. **(B)** SEC of the purified ^ZIP-MBP^Mga2^950-1062^ protein in the detergent-containing SEC-buffer. The protein concentration was adjusted to the indicated concentrations, and 100 *μ*l of each of these samples were subjected to SEC using a Superdex 200 10/300 Increase column. **(C)** SEC of the purified non-zipped ^MBP^Mga2^950-1062^ protein in SEC-buffer. The protein concentration was adjusted to the indicated concentrations, and 100 *μ*l of each of these sampleswere loaded onto a Superdex 200 10/300 Increase column. **(D)** Sucrose-density gradients centrifugation for proteoliposomes containing ^ZIP-MBP^Mga2^950-1062^ at a molar protein:lipid ratio of 1:8,000. The proteoliposome sample was adjusted to 40% w/v sucrose and overlaid with sucrose cushions of different concentrations (20%, 10%, 5%, 0% w/v). After ultracentrifugation, 13 fractions were collected from top to bottom. The relative content of lipids in the individual fractions was determined by Hoechst 33342 fluorescent staining. The amount of ^MBP^Mga2-TMH in the fractions was monitored by immunoblotting using anti-MBP antibodies. **(E)** *In vitro* ubiquitylation reactions were performed with the WT ^ZIP-MBP^Mga2^950-1062^ sense-and-response construct reconstituted in the indicated lipid environments at a protein:lipid ratio of 1:8,000. After indicated times, the reactions were stopped and subjected to SDS-PAGE. For analysis, an immunoblot using anti-MBP antibodies was performed.

**Figure S2.**
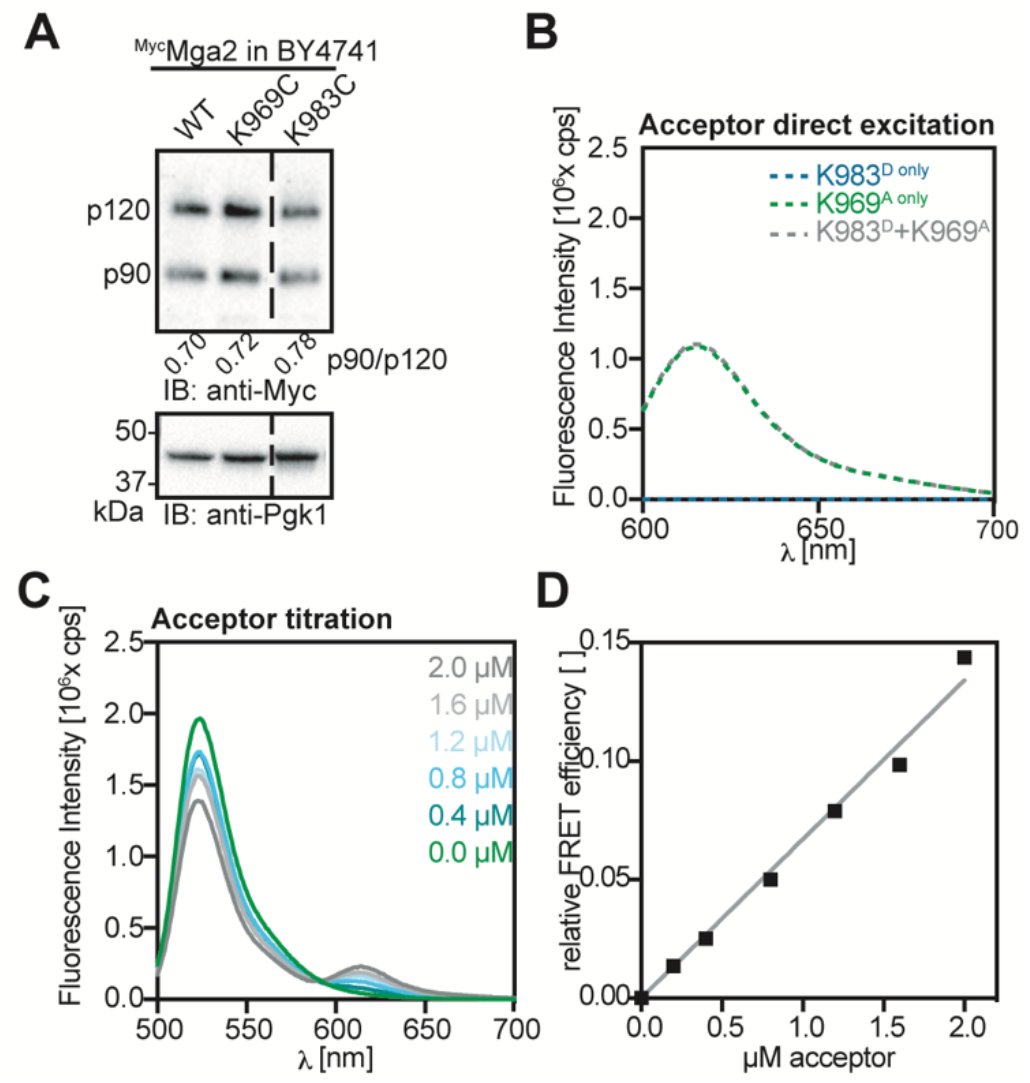
Establishing a FRET reporter based on sense-and-respond construct. **(A)** Immunoblot analysis of indicated ^Myc^Mga2 variants produced at near-endogenous levels in the BY4741 wild type background. Cells were cultivated in YPD to the mid-logarithmic growth phase. Crude cell lysates were subjected to SDS-PAGE and analyzed by immunoblotting using anti-Myc antibodies. The Mga2 p90:p120 ratios were determined by densiometric quantification using Fiji. An anti-Pgk1 immunoblot served as loading control. **(B)** Fluorescence emission spectra for the samples in shown in Figure 3C upon direct acceptor excitation at 590 nm). **(C)** 2 *μ*M donor was titrated with the indicated acceptor concentrations and fluorescence emission spectra were measured upon donor excitation. The overall protein concentrations were maintained by the use of unlabeled ^ZIP-MBP^Mga2^950-1062^. **(D)** Relative FRET efficiencies were determined from the donor/acceptor intensity ratios in (C). Data were fitted via linear regression.

**Figure S3.**
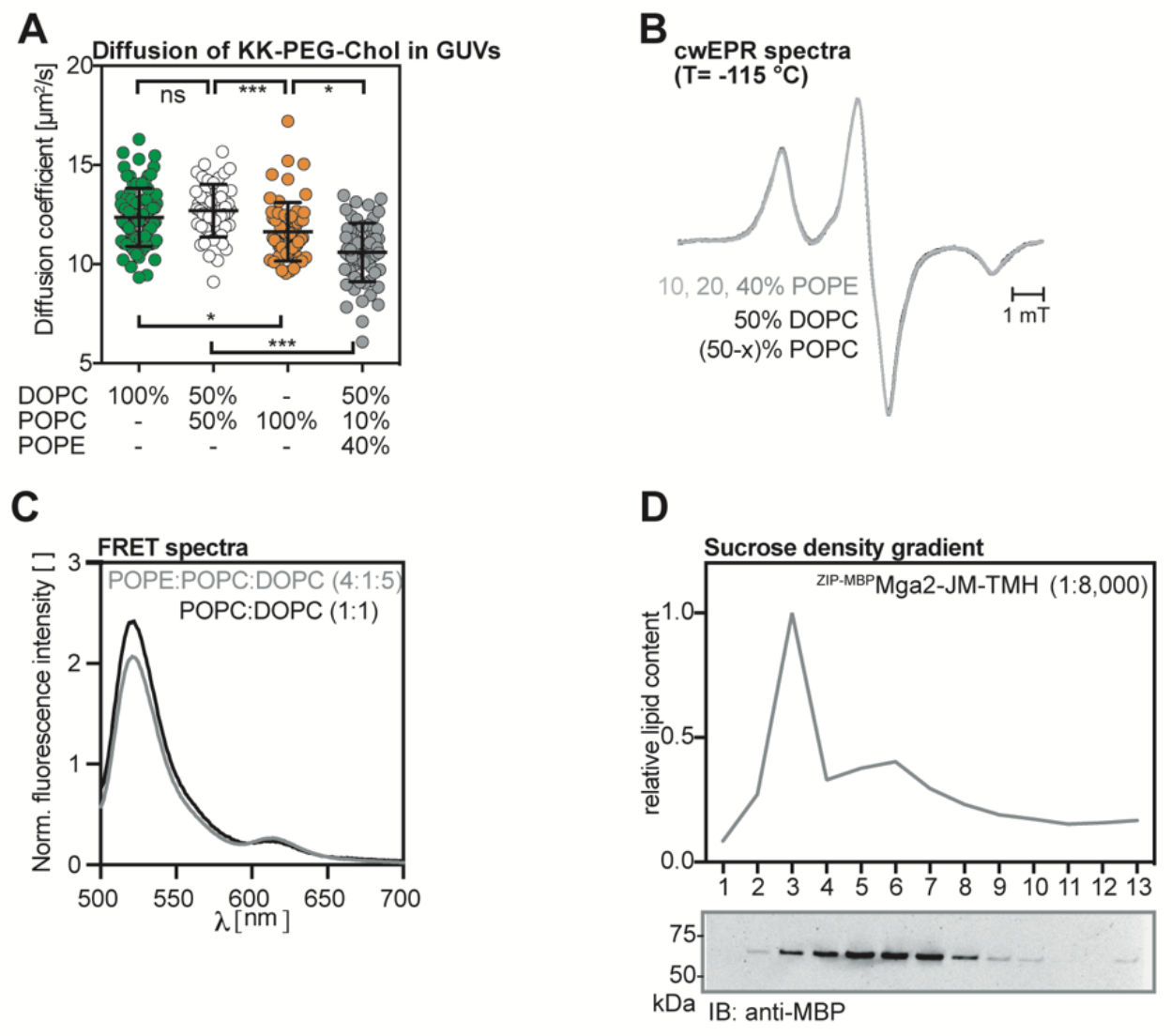
Reconstituting sense-and-response construct in PE-containing liposomes. **(A)** Diffusion coefficients of Star Red-PEG Cholesterol in giant unilaminar vesicles of the indicated lipids were determined by confocal point-FCS. Plotted Is the mean ± SD (n ≥ 55). A Kolmogorov-Smirnov test was performed to test for statistical significance (*p<0.05, ***p<0.001). **B)** Intensity normalized cwEPR spectra recorded at −115°C for a fusion protein composed of MBP and the TMH of Mga2 (^MBP^Mga2^1032-1062^) labeled at position W1042C was reconstituted at a molar protein:lipid of 1:500 in liposomes composed of the indicated lipid mixtures. **(C)** Fluorescence emission spectra of the (K983^D^+K969^A^) FRET pair reconstituted in liposomes composed of the indicated lipid mixtures were recorded (ex: 488 nm, em: 500-700 nm), normalized to the maximal acceptor emission after direct acceptor excitation (ex: 590 nm), and plotted. The emission spectra were normalized to acceptor emission after direct acceptor excitation. **D)** Sucrose-density gradient centrifugation for proteoliposomes containing ^ZIP-MBP^Mga2^950-1062^ at a molar protein:lipid ratio of 1:8,000 in a lipid mixture of 50 mol% DOPC, 10 mol% POPC and 40 mol% POPE. Samples were adjusted to 40% sucrose and overlaid with decreasing concentrations of sucrose-solution (20%, 10%, 5%, 0%). After ultracentrifugation 13 fractions were recovered from from top to bottom. The relative content of lipids in the individual fractions was determined by Hoechst 33342 fluorescent staining. The amount of ^MBP^Mga2-TMH in the fractions was monitored by immunoblotting using anti-MBP antibodies.

**Figure S4.**
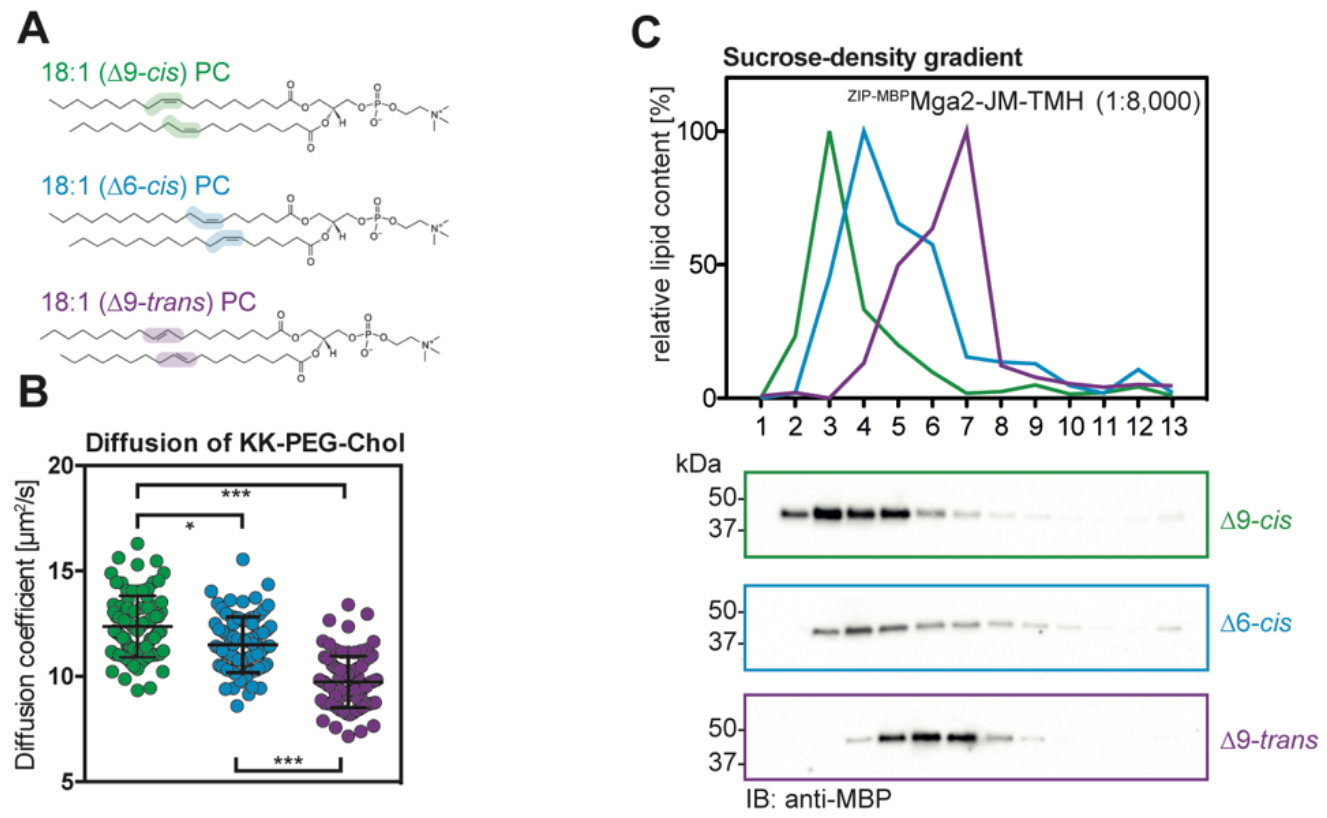
Reconstituting sense-and-response construct in liposomes with different PC-species. **(A)** Chemical structure of the three relevant PC lipids with distinct double bonds isomers and positions. All lipids contain a PC head group, two acyl chains of 18 carbons with one double bond. They only differ in the position (Δ9 or Δ6) and the orientation of the double bond (*cis* or *trans*). The color code is maintained in (B, C). (Structures adapted from avantilipids.com) **(B)** Diffusion coefficients of Star Red-PEG Cholesterol in giant unilaminar vesicles of the indicated lipids were determined by confocal point FCS. Plotted is the mean ± SD (n ≥ 85). A Kolmogorov-Smirnov test was performed to test for statistical significance (*p<0.05, ***p<0.001) **(C)** Sucrose-density gradient centrifugation for proteoliposomes containing ^ZIP-MBP^Mga2^950-1062^ at a molar protein:lipid ratio of 1:8,000 prepared with the indicated lipids. Samples were adjusted to 40% sucrose and overlaid with decreasing concentrations of sucrose-solution (20%, 10%, 5%, 0%). After ultracentrifugation fractions were taken off from top to bottom. The relative lipid content of the individual fractions was determined by Hoechst 33342 fluorescent staining. The amount of ^MBP^Mga2-TMH in the fractions was monitored by immunoblotting using anti-MBP antibodies.

**Figure S5.**
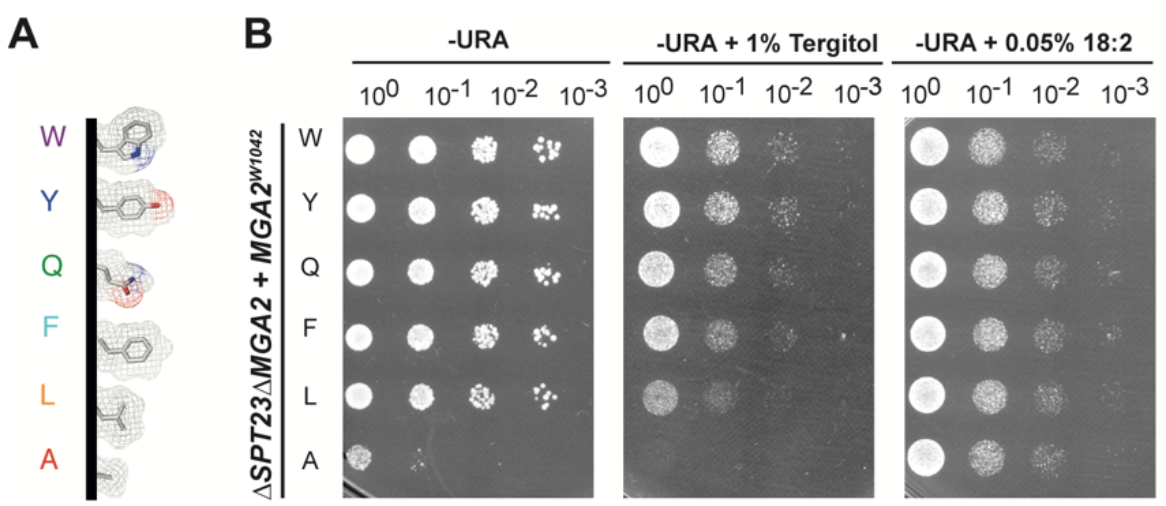
Mutagenesis of sensory residue W1042 and phenotypic charaterization. **(A)** Representations of the amino acids (and substitutions) at position of the sensory W1042 in the TMH of Mga2. The side-chain structures were modeled using PyMOL and are shown as sticks with electron meshes. **(B)** Spotting test for rescue of UFA auxotrophy. The indicated *MGA2* variants were expressed from their endogenous promoters on *CEN*-based plasmids in the Δ*SPT23*Δ*MGA2* strain background. Cultures were depleted for UFA for 5 h before serial dilutions in SCD-URA (10^0^, 10^−1^, 10^−2^, 10^−3^) were prepared and spotted onto SCD-URA plates supplemented with the indicated additives. Colony growth was documented after 2 days incubation at 30 °C.

**Table S1.** Plasmids used in this study.

| Plasmid | Description | Source |
| --- | --- | --- |
| *in vivo* |  |  |
| pRS316 | Empty vector (CEN6-ARS4, URA3, AMP) | EUROSCARF |
| pRE262 | pRS316-3xMyc-*MGA2* WT | 26 |
| pRE266 | pRS316-3xMyc-*MGA2* W1042A | 26 |
| pRE305 | pRS316-3xMyc-*MGA2* W1042L | 26 |
| pRE333 | pRS316-3xMyc-*MGA2* W1042Y | This study |
| pRE334 | pRS316-3xMyc-*MGA2* W1042F | This study |
| pRE335 | pRS316-3xMyc-*MGA2* W1042Q | This study |
| pRE683 | pRS316-3xMyc-*MGA2* K969C | This study |
| pRE684 | pRS316-3xMyc-*MGA2* K983C | This study |
| *in vitro* |  |  |
| pRE345 | pMALC-2x-MBP-*MGA2*-TMH W1042C | 26 |
| pRE496 | pETM-m60-8xHis-hUb WT | This study (kindly provided by V. Dötsch) |
| pSB125 | pMALC-2x-MBP-*MGA2*-JM-TMH WT | This study |
| pSB174 | pMALC-2x-ZIP-MBP-*MGA2*-JM-TMH WT | This study |
| pSB181 | pMALC-2x-ZIP-MBP-*MGA2*-JM-TMH ΔLPKY | This study |
| pSB182 | pMALC-2x-ZIP-MBP-*MGA2*-JM-TMH W1042A | This study |
| pSB186 | pMALC-2x-ZIP-MBP-*MGA2*-JM-TMH K980R, K983R, K985R | This study |
| pSB187 | pMALC-2x-ZIP-MBP-*MGA2*-JM-TMH K983C | This study |
| pSB188 | pMALC-2x-ZIP-MBP-*MGA2*-JM-TMH K969C | This study |
| pSB189 | pMALC-2x-ZIP-MBP-*MGA2*-JM-TMH K969C, W1042A | This study |
| pSB190 | pMALC-2x-ZIP-MBP-*MGA2*-JM-TMH K983C, W1042A | This study |

**Table S2.** Strains used in this study.

| Strain Number | Description | Genotype | Source | Plasmid |
| --- | --- | --- | --- | --- |
| YRE001 | BY4741 | MATa; his3Δ1; leu2Δ0; met15Δ0; ura3Δ0 | EUROSCARF <sup>64</sup> |  |
| YRE009 | *ΔUBX2* | MATa; his3Δ1; leu2Δ0; met15Δ0; ura3Δ0; ubx2Δ::kanMX4 | EUROSCARF <sup>65</sup> |  |
| YRE067 | BY4741 3xMyc-*MGA2* WT | MATa; his3Δ1; leu2Δ0; met15Δ0; ura3Δ0 | <sup>26</sup> | pRE262 |
| YRE068 | BY4741 3xMyc-*MGA2* W1042A | MATa; his3Δ1; leu2Δ0; met15Δ0; ura3Δ0 | <sup>26</sup> | pRE266 |
| YRE071 | *ΔUBX2* 3xMyc-*MGA2* WT | MATa; his3Δ1; leu2Δ0; met15Δ0; ura3Δ0; ubx2Δ::kanMX4 | <sup>26</sup> | pRE262 |
| YRE199 | BY4741 3xMyc-*MGA2* W1042L | MATa; his3Δ1; leu2Δ0; met15Δ0; ura3Δ0 | <sup>26</sup> | pRE305 |
| YRE216 | BY4741 3xMyc-*MGA2* W1042Y | MATa; his3Δ1; leu2Δ0; met15Δ0; ura3Δ0 | This study | pRE333 |
| YRE217 | BY4741 3xMyc-*MGA2* W1042F | MATa; his3Δ1; leu2Δ0; met15Δ0; ura3Δ0 | This study | pRE334 |
| YRE228 | *ΔSPT23, ΔMGA2* | MATα; his3Δ1; leu2Δ0; lys2Δ0; ura3Δ0; spt23Δ::kanMX4; mga2Δ::natMX | Kindly provided by Harald Hofbauer (Graz) |  |
| YRE295 | *ΔSPT23, ΔMGA2* 3xMyc-*MGA2* WT | MATα; his3Δ1; leu2Δ0; lys2Δ0; ura3Δ0; spt23Δ::kanMX4; mga2Δ::natMX | This study | pRE262 |
| YRE296 | *ΔSPT23, ΔMGA2* 3xMyc-*MGA2* W1042A | MATα; his3Δ1; leu2Δ0; lys2Δ0; ura3Δ0; spt23Δ::kanMX4; mga2Δ::natMX | This study | pRE266 |
| YRE297 | *ΔSPT23, ΔMGA2* 3xMyc-*MGA2* W1042L | MATα; his3Δ1; leu2Δ0; lys2Δ0; ura3Δ0; spt23Δ::kanMX4; mga2Δ::natMX | This study | pRE305 |
| YRE404 | BY4741 3xMyc-*MGA2* W1042Q | MATa; his3Δ1; leu2Δ0; met15Δ0; ura3Δ0 | This study | pRE335 |
| YRE415 | BY4741 empty vector pRS316 | MATa; his3Δ1; leu2Δ0; met15Δ0; ura3Δ0 | This study | pRS316 |
| YRE572 | *ΔSPT23, ΔMGA2* 3xMyc-*MGA2* W1042Q | MATα; his3Δ1; leu2Δ0; lys2Δ0; ura3Δ0; spt23Δ::kanMX4; mga2Δ::natMX | This study | pRE335 |
| YRE573 | *ΔSPT23, ΔMGA2* 3xMyc-*MGA2* W1042F | MATα; his3Δ1; leu2Δ0; lys2Δ0; ura3Δ0; spt23Δ::kanMX4; mga2Δ::natMX | This study | pRE334 |
| YRE574 | *ΔSPT23, ΔMGA2* 3xMyc-*MGA2* W1042Y | MATα; his3Δ1; leu2Δ0; lys2Δ0; ura3Δ0; spt23Δ::kanMX4; mga2Δ::natMX | This study | pRE333 |
| YRE578 | *ΔSPT23, ΔMGA2* empty vector pRS316 | MATα; his3Δ1; leu2Δ0; lys2Δ0; ura3Δ0; spt23Δ::kanMX4; mga2Δ::natMX | This study | pRS316 |

